# Revisiting the impact of synthetic ORF sequences on engineered LINE-1 retrotransposition

**DOI:** 10.1101/2022.08.29.505632

**Authors:** Dorothy Chan, Stephanie Workman, Nathan Smits, Alexander L. Carleton, Patricia Gerdes, Jeffrey S Han, Jef D Boeke, Geoffrey J Faulkner, Sandra R Richardson

## Abstract

The retrotransposon Long Interspersed Element-1 (L1) contains adenosine rich open reading frames (ORFs), a characteristic that limits its expression in mammalian cells. A synthetic mouse L1 (smL1) with ORF adenosine content decreased from 40% to 26% showed increased mRNA expression and retrotransposed far more efficiently than the native parental element, L1spa (1). Here, we observe two nonsynonymous substitutions between the L1spa and smL1 ORF1 sequences, and note that the smL1 3’UTR lacks a conserved guanosine-rich region (GRR) which could take on a G-quadruplex secondary structure. We find that the combined effect of the altered ORF1p amino acid sequence and the GRR 3’UTR deletion, rather than synthetic ORF sequences, accounts for the increase in smL1 retrotransposition efficiency over L1spa. Furthermore, we demonstrate that the presence and position of the GRR within the L1 reporter construct impact mouse L1 ORF1p expression and retrotransposition efficiency. Our results prompt a reevaluation of synthetic L1 activity and suggest that the manner in which L1 sequences are cloned into engineered reporter vectors has, in some cases, resulted in an underestimation of native mouse L1 retrotransposition efficiency.

**Author Summary:** L1 retrotransposons are mobile DNA elements or “jumping genes” that can copy-and-paste their sequences to new locations in the host genome. The jumping ability, or retrotransposition efficiency, of individual L1 elements can be evaluated using a cultured cell assay in which the L1 is tagged in its 3’ untranslated region (3’UTR) with a reporter gene that becomes expressed upon successful retrotransposition. In a previous study, authors Han and Boeke reported that the retrotransposition efficiency of a mouse L1 element could be enhanced dramatically by synthetically increasing the GC content of the L1 open reading frames (ORFs) without changing their amino acid sequence. Curiously, a similarly constructed synthetic human L1 achieved only a modest increase in retrotransposition efficiency over the native element. Here, we find that two coding changes and a partial deletion comprising a guanine-rich region (GRR) of the mouse L1 3’UTR sequence which occurred during construction of the synthetic mouse L1 reporter are responsible for the increased jumping of the synthetic mouse L1 construct relative to the native L1spa element. We find that the presence/absence and also the placement of this GRR 3’UTR region within the reporter construct impact ORF1p expression and engineered L1 retrotransposition efficiency. Together, our study reconciles the disparate impacts of synthetic sequences upon human and mouse L1 retrotransposition efficiency, prompts a reconsideration of numerous studies using synthetic L1 constructs, and will inform the ongoing use of synthetic and natural mouse L1 reporter constructs in vivo and in vitro.

## Introduction

L1 retrotransposons are an ongoing source of mutagenesis in mammalian genomes (2, 3). Mice contain ∼3000 retrotransposition-competent L1s (RC-L1s) representing three subfamilies (L1_T_F_, L1_G_F_, L1_A), distinguished by their monomeric promoter-harboring 5’UTRs (4–9). RC-L1s encode two proteins required for their mobility: ORF1p, a nucleic acid binding and chaperone protein, and ORF2p, which has endonuclease and reverse transcriptase activities critical for the generation of new L1 insertions by target-primed reverse transcription (TPRT) (10–19). The L1 3’UTR contains a weak polyadenylation (poly-A) signal and a poly-purine tract that is found in L1s across species which varies in length and nucleotide sequence (20–24). Hallmarks of TPRT-mediated L1 integrants include insertion at the degenerate consensus sequence 5’-TTTTT/AA-3’, 3’ terminal poly-A tracts, and flanking target site duplications (TSDs)(25–30). Approximately 1 in 8 mice and 1 in 62.5 humans harbor a *de novo* L1 insertion (31, 32). Moreover, >100 cases of human genetic disease and numerous spontaneous mouse mutants have been linked to L1-mediated retrotransposition events, highlighting the impact of L1 mutagenesis on mammalian genomes (33, 34).

The retrotransposition efficiency of individual L1 copies can be evaluated in cultured cells and *in vivo* (19, 35–37). In these assays, a retrotransposition indicator cassette (4), consisting of a reporter gene in antisense orientation to the L1 and equipped with its own promoter and polyadenylation (poly-A) signal, is placed into the L1 3’UTR (Figure 1A). The reporter gene is interrupted by an intron oriented in sense to the L1, with expression achieved only upon splicing and reverse transcription of the L1 reporter mRNA to deliver an intact copy of the reporter cassette into genomic DNA. Expression and polyadenylation of the full-length L1 mRNA can be augmented by a strong heterologous promoter and poly-A signal flanking the L1 element and reporter cassette (19, 38). In selection-based cultured cell assays, quantification of antibiotic-resistant foci provides a readout of retrotransposition efficiency (19).

**Figure 1.**
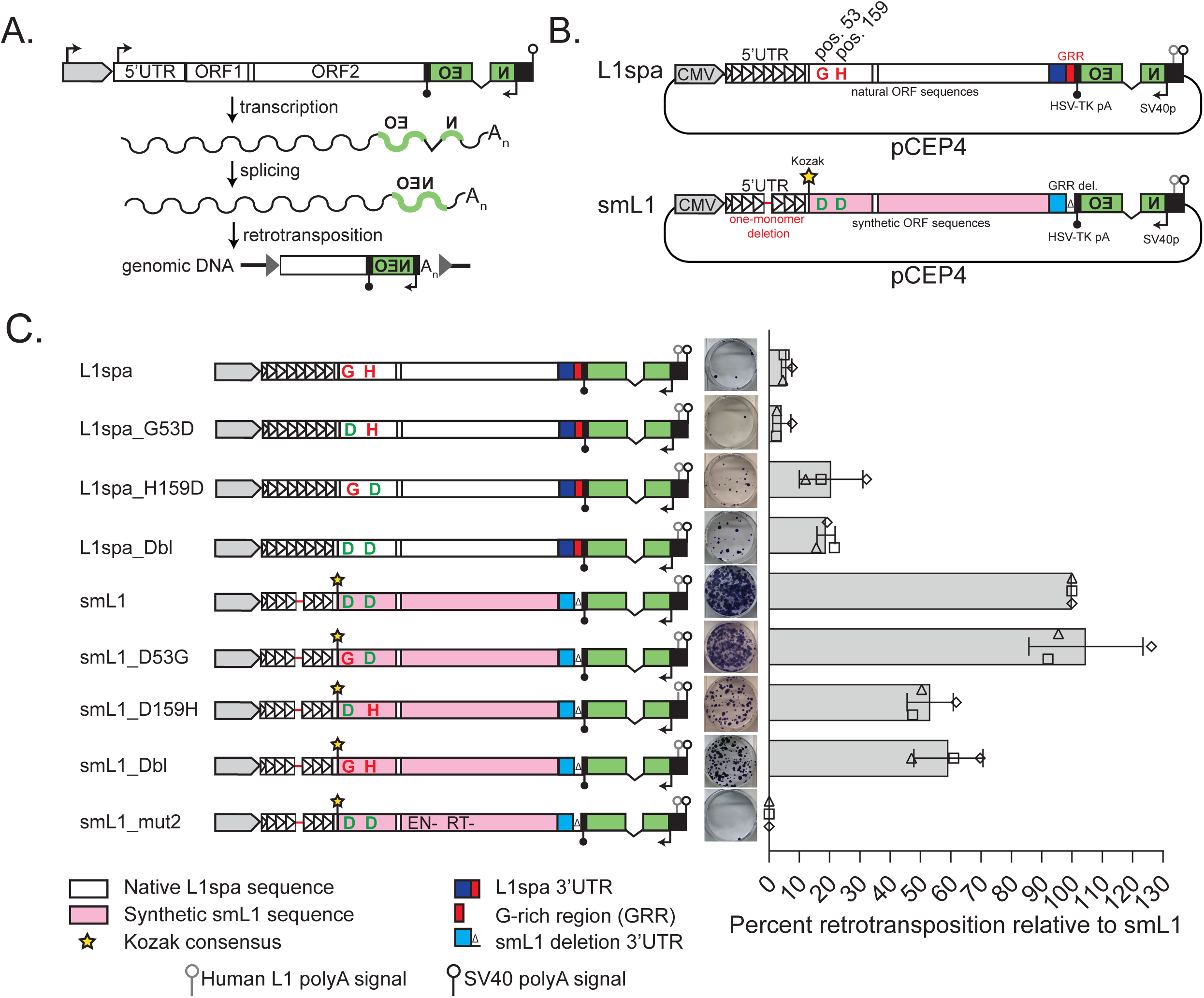
L1spa and smL1 differ at two ORF1p amino acid positions. A. The cultured cell retrotransposition assay. A full-length L1 containing intact open reading frames (ORF1 and ORF2) is driven by its native 5’UTR promoter or a strong heterologous promoter (grey) and equipped with a polyadenylation signal (native and/ or engineered, open lollipop). The L1 is tagged with a retrotransposition indicator cassette (green) inserted in its 3’UTR. The reporter consists of a backwards intron-containing neomycin phosphotransferase (NEO) reporter gene in opposite transcriptional orientation to the L1 and equipped with an SV40 promoter and HSV-TK polyadenylation signal (upside-down black arrow and black lollipop, respectively). Expression of the reporter gene from genomic DNA is achieved upon retrotransposition. B. L1spa/pTN201 and smL1 plasmids. Features are depicted as follows: CMV promoter (grey), L1spa 5’UTR (white triangles, representing monomers), smL1 5’UTR (one-monomer deletion represented by red line). NEO reporter cassette (green) with SV40 early promoter (black arrow) and HSV-Tk polyadenylation signal (filled black lollipop). Native L1spa nucleotide sequence (white); synthetic sequence (pink). The amino acids at positions 53 and 159 in ORF1p are indicated. Engineered Kozak consensus sequence (star). L1spa 3’UTR (dark blue); G-rich region (red). Mutated smL1 3’UTR (light blue), deleted GRR (Δ symbol). Human L1 polyadenylation signal (residual from original human retrotransposition indicator constructs (6, 19); open grey lollipop) and the SV40 polyadenylation signal (open black lollipop). C. Effects of ORF1p amino acid changes on retrotransposition. From left to right: a schematic of each construct (as described in B), a representative well from the retrotransposition assay, and its retrotransposition efficiency relative to smL1. The histogram displays the mean of three biological replicate assays, each comprising three technical replicates per construct. Triangle, square, and diamond shapes indicate the individual values for each biological replicate, and the error bars represent standard deviation among biological replicates.

The adenosine richness of L1 ORFs limits L1 expression in mammalian cells and influences expression of genes containing intronic sense oriented L1 insertions (39–41). A synthetic mouse L1 sequence (smL1, or ORFeus_Mm) previously was developed by Boeke and Han to increase the retrotransposition efficiency of engineered L1 reporter constructs, improving their utility for random mutagenesis screens of mammalian genomes (1). smL1 was derived from the L1_T_F_ subfamily element L1spa (6, 42) whereby the adenosine content of the L1spa ORFs was reduced from 40% to 26% via synonymous substitutions. The synthetic ORF sequences of smL1 yielded markedly increased mRNA expression levels, and smL1 retrotransposed ∼200-fold more efficiently in HeLa cells than the retrotransposition indicator construct pTN201/L1spa (1, 6). Notably, similar constructs containing a synthetic human L1 showed at most a 3-fold increase in retrotransposition efficiency over the native element (43). Here, we note both coding and non-coding differences between smL1 and pTN201/L1spa, and we systematically evaluate the impact of these differences, and of synthetic ORF sequences, on mouse L1 retrotransposition efficiency. We determine that the presence and position of a conserved poly purine tract in the mouse L1 3’UTR profoundly impacts the retrotransposition efficiency of engineered mouse L1 reporter constructs and investigate potential mechanisms by which this impact may arise.

## Results

We sequenced the entire pCEP4-based smL1 and pTN201/L1spa plasmids by capillary sequencing and Oxford Nanopore Technologies (ONT) long-read DNA sequencing (Supplemental Table 1) and observed that the synthetic sequences of the smL1 ORFs largely comprised synonymous nucleotide substitutions relative to L1spa. However, smL1 ORF1 contained two nonsynonymous substitutions (G53D and H159D) and bore an engineered Kozak consensus at its initiation codon that was absent from pTN201/L1spa. Furthermore, the terminal 159 bp of 3’UTR sequence present in pTN201/L1spa construct was absent from smL1 (Figure 1B; Supplemental Figure 1). We refer to this 159 bp sequence as the guanine-rich region (GRR), as it encompasses the conserved poly-purine tract. We traced this deletion to the inadvertent loss of a NdeI restriction fragment during subcloning of the smL1 construct (Supplemental Figure 2). In addition, in the smL1 3’UTR nucleotides 2-18, encompassing a SacI restriction site, have been replaced with an 11 nucleotide sequence including an engineered AgeI restriction site, and an engineered PacI site in the inter-ORF spacer sequence introduces three nucleotide changes. The remaining smL1 3’UTR contains 19 nucleotide substitutions and two single nucleotide deletions relative to the L1spa 3’UTR, which arose during synthesis of the smL1 construct (Supplemental Figure 1). Long-read sequencing also revealed the deletion of one monomer unit from the 5’UTR promoter of the smL1 construct relative to the pTN201/L1spa construct (Figure 1B).

The L1_T_F_ subfamily consensus, and previously described active mouse L1_T_F_ elements, contain aspartic acid at ORF1p positions 53 and 159, rather than glycine and histidine respectively, as found in L1spa ORF1p (6, 8, 44, 45). Indeed, the substitution H159D improves L1spa ORF1p nucleic acid chaperone activity and retrotransposition efficiency without affecting steady-state ORF1p levels (46, 47). To quantify the contribution of G53D and H159D to the increased retrotransposition efficiency of smL1 over L1spa, we generated L1spa_G53D, L1spa_H159D, and a construct with both changes, L1spa_Dbl (Figure 1C). In a HeLa cell transient transfection-based retrotransposition assay (35, 37), L1spa mobilized at ∼6% the efficiency of smL1, and, consistent with previous results (46), the substitution G53D had little effect on retrotransposition. However, L1spa_H159D and L1spa_Dbl mobilized at ∼19% the efficiency of smL1. We likewise generated smL1_D53G, smL1_D159H, and smL1_Dbl (D53G and D159H), and found that while smL1_D53G retrotransposed with similar efficiency to smL1, smL1_D159H and smL1_Dbl exhibited a ∼40% reduction in retrotransposition efficiency relative to smL1 (Figure 1C). As expected, the negative control construct smL1_mut2, which has inactivating mutations in both the endonuclease and reverse transcriptase domains of ORF2 (1), did not retrotranspose. Thus, the H159D amino acid substitution partially explains the increased retrotransposition efficiency of smL1 over L1spa. The G53 and H159 variants apparently are restricted to L1spa and closely related elements, as the L1_TF consensus contains both D53 and D159 (6–8, 46–48). As L1spa_Dbl and smL1 have identical ORF1p amino acid sequences, we used L1spa_Dbl rather than L1spa_H159D as the basis for comparison to smL1-derived constructs in subsequent experiments.

To determine the contribution of non-coding changes between smL1 and L1spa_Dbl to their differing retrotransposition efficiencies, we equipped L1spa_Dbl ORF1 with a Kozak consensus and replaced the L1spa 3’UTR with the deletion-containing smL1 3’UTR to generate “L1spa Double in smL1 backbone”, hereafter abbreviated L1spa_Dbl_smBB. Despite entirely lacking synthetic ORF sequences, L1spa_Dbl_smBB retrotransposed with equal efficiency to smL1 (Figure 2A). When tested independently, removing the Kozak consensus from this construct (L1spa_Dbl_smBB_Delete_Kozak) had no impact on retrotransposition efficiency. However, restoring the GRR-containing L1spa 3’UTR (L1spa_Dbl_smBB_restore_3’UTR) reduced retrotransposition to similar levels as L1spa_Dbl (Figure 2A). This result suggested that the deletion-containing smL1 3’UTR largely accounts for the increased retrotransposition efficiency of L1spa_Dbl_SMBB over L1spa. To determine the impact of the GRR in the context of a synthetic L1 element, we generated smL1_spa3’UTR, which contains synthetic smL1 ORF sequences and the full GRR-bearing L1spa 3’UTR. Surprisingly, replacing the GRR into smL1 did not diminish retrotransposition relative to the original smL1 construct (Figure 2A). Taken together, our results indicate that the previously reported improvement in smL1 retrotransposition over L1spa is not attributable to the increased GC content of the smL1 ORFs, but to the combined deleterious effect of the H159 substitution and inclusion of the GRR upstream of the NEO cassette in the pTN201/L1spa construct. While the presence of a histidine residue rather than aspartic acid at ORF1p position 159 diminishes retrotransposition of both L1spa and smL1 (Figure 1C), retrotransposition of the synthetic mouse L1 was not affected by inclusion of the GRR upstream of the NEO cassette.

**Figure 2.**
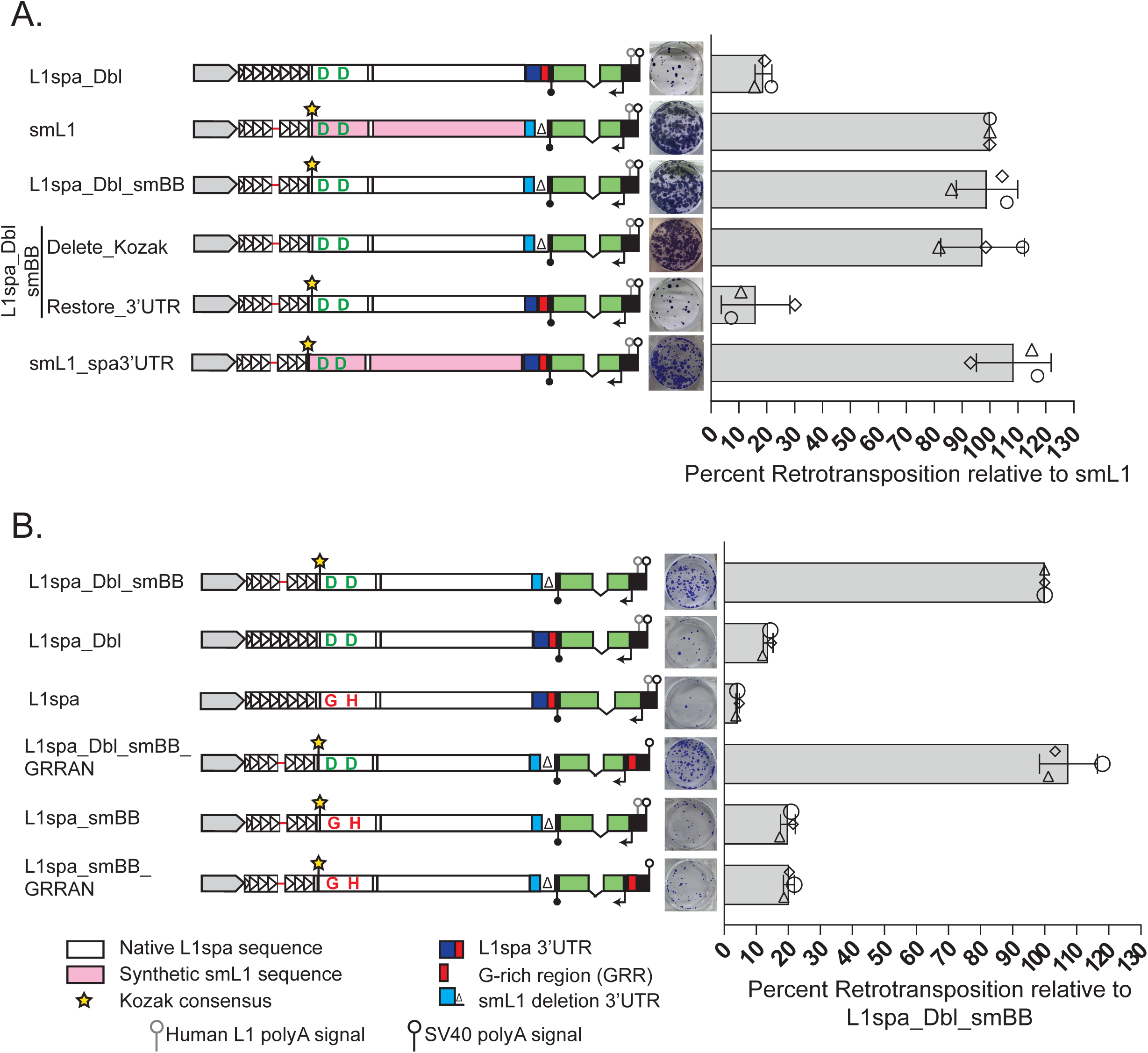
The presence and position of the L1spa 3’UTR G-rich region impact retrotransposition. A. From left to right: a schematic of each construct, a representative well from the retrotransposition assay, and its retrotransposition efficiency relative to smL1. L1spa_Dbl_smBB contains the ORF1 and ORF2 sequences of L1spa_Double with a Kozak consensus at the ORF1p initiation codon (as in smL1), and the deletion 3’UTR of the smL1 construct. Delete_Kozak is equivalent to L1spa_Dbl_smBB but with the Kozak sequence removed. Restore_3’UTR is equivalent to L1spa_Dbl_smBB but with the full L1spa 3’UTR restored upstream of the NEO cassette. smL1_spa3’UTR is equivalent to Restore_3’UTR but with the smL1 ORFs. Each histogram displays the mean of three biological replicate assays, each comprising three technical replicates per construct. Triangle, square, and diamond shapes indicate the individual values for each biological replicate, and the error bars represent standard deviation among biological replicates. B. As above; L1spa_Dbl_smBB_GRRAN is equivalent to L1spa_Dbl_smBB but with the G-rich region of the L1spa 3’UTR placed downstream of the NEO cassette, and the human L1 polyadenylation signal removed. L1spa_SMBB is equivalent to L1spa_Dbl_SMBB but with the original L1spa (G53, H159) ORF1p sequence. L1spa_SMBB_GRRAN is equivalent to L1spa_Dbl_SMBB_GRRAN but with the original L1spa ORF1p sequence. Each histogram displays the mean of three biological replicate assays, each comprising three technical replicates per construct. Triangle, square, and diamond shapes indicate the individual values for each biological replicate, and the error bars represent standard deviation among biological replicates.

We next investigated the sequence characteristics of the L1spa GRR responsible for diminished L1spa retrotransposition efficiency. To test whether the non-GRR portion of the L1spa 3’UTR sequence impacts retrotransposition, we generated L1spa_Dbl_smBB_Restore_L1spa_nonGRR (Supplemental Figure 3A). This construct retrotransposed similarly to L1spa_Dbl_smBB, indicating that the non-GRR portion of the L1spa 3’UTR by itself has little impact on retrotransposition efficiency (Supplemental Figure 3A). In contrast, an L1spa_Dbl construct containing the L1spa GRR and lacking the remaining 3’UTR sequence retrotransposed comparably to L1spa_Dbl, confirming that the L1spa GRR alone is sufficient to impede retrotransposition of a natural mouse L1 (L1spa_Dbl_smBB_Restore_L1spa_GRR, Supplemental Figure 3A). To determine whether the negative impact on engineered retrotransposition is common to the GRR sequences from ancient (V, F) and presently active (G_F_, A) L1s in addition to the T_F_ subfamily (represented by L1spa), we replaced the L1spa GRR with the consensus GRR sequence from each of these subfamilies, which contain 9 (G_F_, A) or 10 (V, F) runs of 3 or more guanine residues (8). In all cases, retrotransposition was reduced to <20% of L1spa_Dbl_smBB (Supplemental Figure 3A). To examine in greater detail the role of the L1spa GRR sequence in retrotransposition inhibition, we generated a series of deletion constructs. The L1spa GRR contains 9 runs of 3 or more guanine residues, separated by variable stretches of other bases (Supplemental Figure 3B). Relative to L1spa_Dbl_SMBB (no GRR sequence), the sequential restoration of guanine runs in a 5’-to-3’ direction resulted in a progressive diminution of retrotransposition efficiency, with the largest difference observed between the constructs containing 5 and 6 guanine runs (Supplemental Figure 3C). Thus, the inhibitory effect of the GRR on engineered L1 retrotransposition is shared by both ancient and presently active natural mouse L1s, and the magnitude of inhibition is positively correlated with the length of the GRR.

Previous studies have reported efficient retrotransposition of natural mouse L1 reporter constructs engineered so that the GRR is positioned downstream of the NEO cassette (46, 47). To evaluate directly how placement of the GRR relative to the NEO cassette impacts retrotransposition efficiency of a natural mouse L1, we placed the L1spa GRR into L1spa_Dbl_SMBB downstream of NEO but upstream of the pCEP4 vector SV40 polyadenylation signal, to generate L1spa_Dbl_smBB_GRR_after_NEO, hereafter abbreviated as L1spa_Dbl_SMBB_GRRAN. This construct retrotransposed with equal efficiency to L1spa_Dbl_smBB (Figure 2B). This result suggests that the impact of the GRR upon engineered natural mouse L1 retrotransposition efficiency is position dependent. We also asked whether removing or relocating the GRR can enable efficient retrotransposition of the original L1spa element. We generated a construct containing the original L1spa ORF1p sequence (G53 and H159) but with the GRR deleted (L1spa_SMBB) or placed downstream of the NEO cassette (L1spa_SMBB_GRR_after_NEO). Both constructs retrotransposed at ∼20% the efficiency of L1spa_Dbl_smBB or L1spa_Dbl_smBB_GRRAN, constituting a slight improvement over L1spa (Figure 2B). We conclude that deletion or repositioning of the GRR cannot compensate for the previously reported (46, 47) effects of H159 upon L1spa ORF1p nucleic acid chaperone activity and retrotransposition efficiency.

The poly-guanine runs comprising the mouse L1 GRR could potentially form a G-quadruplex secondary structure. G-quadruplexes can form in both DNA and RNA and have been implicated in the impediment of RNA polymerase procession (49–51) and, when located in the UTRs of certain genes, can promote the use of alternative polyadenylation sites (52) and effect translational repression (53). We therefore asked whether the mouse L1 GRR can influence the position at which the L1 transcript is polyadenylated. We reasoned that for constructs in which the GRR is located upstream of the NEO cassette, polyadenylation in the vicinity of the GRR would result in an L1 mRNA molecule lacking the NEO cassette, the retrotransposition of which would be undetectable in a selection-based assay. To detect potential premature polyadenylation, we performed 3’RACE assays on RNA extracted from cells transfected with L1spa_Dbl constructs with the GRR upstream of the NEO cassette (L1spa_Dbl and Restore_3’UTR), downstream of the NEO cassette (L1spa_Dbl_smBB_GRRAN), or deleted entirely (L1spa_Dbl_SMBB). We used primers specific to the engineered L1 reporter transcript: one primer located in the SV40 promoter of the antisense NEO cassette to detect mRNAs polyadenylated as directed by the SV40 polyadenylation signal, and another in the non-GRR segment of the L1spa 3’UTR to detect mRNAs prematurely polyadenylated proximal to the GRR. We also used a primer annealing to the 3’ end of the hygromycin resistance gene in the pCEP4 backbone as an internal control (Figure 3A). For all constructs, we detected the expected ∼520bp band corresponding to polyadenylated transcripts from the hygromycin resistance gene. Using a primer to the antisense SV40 promoter responsible for NEO expression, we detected the expected ∼350 bp band corresponding to L1 reporter transcripts polyadenylated at the SV40 polyadenylation signal for L1spa_Dbl, L1spa_Dbl_SMBB, and Restore_3’UTR, and a ∼520 bp band from L1spa_Dbl_SMBB_GRR_after_NEO, the increased size of which is accounted for by the inclusion of the GRR sequence between the antisense SV40 promoter and the SV40 polyadenylation signal. Using a primer annealing to the L1spa 3’UTR, we did not detect bands in the range of 200-400 bp indicative of polyadenylation within or proximal to the GRR in either the construct L1spa_Dbl or Restore_3’UTR (Figure 3A). Furthermore, 10/10 fully characterized L1 insertions arising from the L1spa_Dbl_smBB_GRRAN construct bore 3’ poly-A tracts downstream of the GRR as directed by the SV40 poly-A signal, rather than aberrant poly-A tracts located upstream of or within the GRR (Figure 3B). Like smL1 insertions previously characterized in HeLa cells (1), all 10 L1spa_Dbl_smBB_GRRAN insertions bore the hallmarks of *bona fide* retrotransposition events including variable-length TSDs and insertion at sites resembling the L1 EN cleavage consensus motif 5’-TTTTT/AA-3’ (Supplemental Figure 5, Supplemental Table 1)(30). Thus, we conclude that aberrant polyadenylation of sense-strand engineered L1 transcripts is unlikely to be responsible for the reduction in retrotransposition efficiency when the GRR is situated upstream of the NEO cassette.

**Figure 3:**
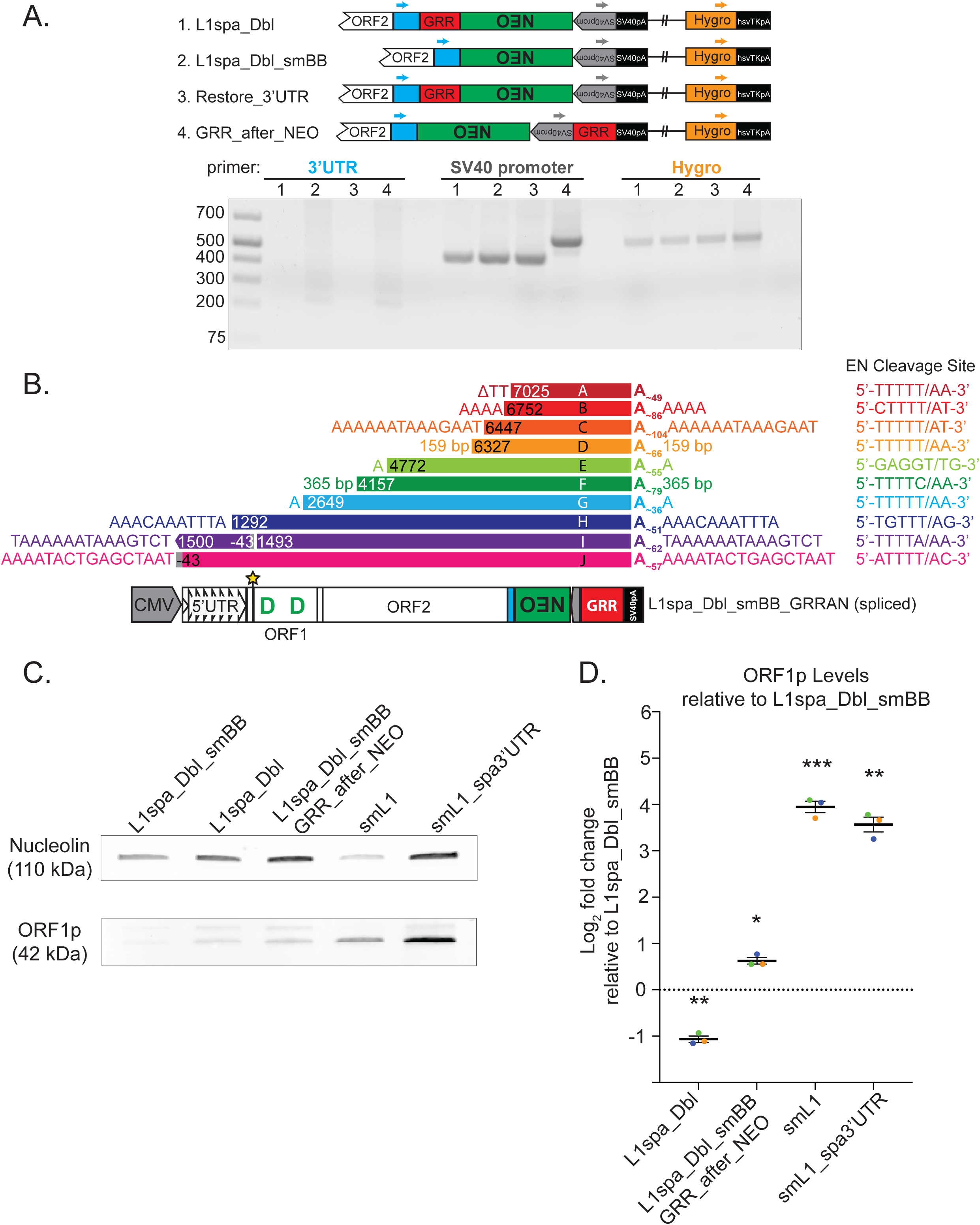
Effects of GRR presence and position on polyadenylation, L1 insertion structure, and ORF1p protein levels. A. 3’RACE assay to identify sites of polyadenylation. Above, schematics representing the constructs 1) L1spa_Dbl, 2) L1spa_Dbl_SMBB, 3) L1spa_Dbl_smBB_Restore_3’UTR and 4) L1spa_Dbl_smBB_GRRAN. The positions of gene-specific primers annealing to the L1spa 3’UTR (blue), the antisense SV40 promoter driving NEO expression (grey), and the hygromycin resistance gene in the pCEP4 vector backbone (orange) are denoted by arrows. Schematics are not drawn to scale. Below, agarose gel showing 3’RACE products detected using RNA isolated from cells transfected with the four above constructs, using each of the indicated primers. B. Structural hallmarks of ten insertions generated with the L1spa_Dbl_smBB_GRR_after_NEO construct are shown, including the 5’ TSD, 5’ truncation position relative to the L1spaDbl_smBB_GRRAN, polyadenylation position, poly-A tail length and 3’TSD; the inferred L1 endonuclease first-strand cleavage site for each insertion is listed to the right. Below, the structure of L1spa_Dbl_smBB_GRRAN, with the spliced NEO cassette in green, the GRR in red, and the SV40 polyadenylation signal in black. Kozak sequence (star shape) and D53, D159 amino acids (green) are indicated. Schematic is not to scale. Delta symbol indicates a target-site deletion. Insertion I contained a 5’ inversion for which the breakpoints are indicated. Insertions I and J arose from a transcript initiating upstream of the L1 5’UTR, at the transcription start site of the CMV promoter within the pCEP4 vector (75) denoted as position -43. For two insertions bearing >100 bp TSDs, the TSD length, rather than TSD sequence, is shown. C. L1 ORF1p expression from reconfigured reporter constructs. Representative Western blot performed on cell extracts from HeLa cells transfected with the indicated reporter construct using a monoclonal antibody specific to mouse L1 ORF1p. Nucleolin was used as a loading control. D. Quantification of nucleolin-normalized ORF1p expression as Log_2_ fold change for L1spa_Dbl, L1spa_Dbl_SMBB_GRR_after_NEO, smL1, and smL1_spa3’UTR relative to L1spa_Dbl_smBB. Each biological replicate (cell lysate from an independent transfection) is represented by a colored dot (rep 1=blue, rep 2=green, rep 3=orange). Horizontal line shows the mean of biological replicates; error bars represent standard error of the mean (SEM). Values for each biological replicate are the mean of two technical replicates (independent blots using the same cell lysate; see Supplemental Figure 6 for individual blots and graphs showing technical replicates contributing to each biological replicate). One sample t and Wilcoxon test; p<0.05*; p<0.01**; p<0.001***.

To investigate the impact of the GRR presence and position upon engineered L1 RNA levels, we performed qRT-PCR on RNA isolated from HeLa cells transfected with the constructs L1spa_Dbl, L1spa_Dbl_SMBB, L1spa_Dbl_SMBB_GRR_after_NEO, smL1, or smL1_spa3’UTR. We employed primer pairs targeting the non-monomeric region of the 5’UTR, the L1spa GRR, and Orf1 and Orf2 sequences of L1spa and smL1. We also targeted a portion of the pCEP4 SV40 polyadenylation signal (SV40pA) that lies upstream of the AAUAAA motif and cleavage site and therefore is expected to be incorporated into the full-length L1 transcriptional unit (Supplemental Figure 4A). The RNA levels for each target were calculated relative to expression of the endogenous housekeeping gene ribosomal protein S18 (*Rps18*).

Across all target sequences, smL1 RNA levels were significantly higher than any of the natural mouse L1 constructs (p<0.05, one-way ANOVA with Tukey’s multiple comparisons test; Supplemental Figure 4, B-E), consistent with previous reports of elevated RNA expression from synthetic L1 constructs (1, 43). Among the natural mouse L1 constructs, RNA levels for L1spa_Dbl_smBB were consistently elevated compared to L1spa_Dbl and L1spa_Dbl_smBB_GRRAN. Although these differences did not reach statistical significance, this result suggests that inclusion of the GRR within the L1 transcriptional unit may reduce L1 RNA levels. Indeed, RNA levels for smL1_spa3’UTR were also diminished relative to smL1 for all target sequences, reaching statistical significance for the Orf2 (p<0.001) and 5’UTR (p<0.05) primer pairs. Primers targeting the GRR sequence itself gave no signal for the GRR-lacking L1spa_Dbl_smBB and smL1 constructs, as expected (Supplemental Figure 4F). GRR RNA levels were lower for L1spa_Dbl compared to L1spa_Dbl_smBB_GRRAN and smL1_spa3’UTR; though not statistically significant, this result suggests that L1spa GRR transcription may be influenced by position within the expression construct or the presence of upstream synthetic ORFs. The above results must be interpreted with the caveat that qRT-PCR signal for short stretches of L1 RNA sequence does not necessarily reflect the abundance of full-length L1 mRNA molecules capable of undergoing retrotransposition. Nevertheless, we conclude that synthetic ORFs and the presence and position of the GRR sequence within the L1 transcriptional unit can modulate engineered L1 RNA levels, but this modulation does not correlate with or explain the observed differences in retrotransposition efficiency (compare Figure 2 and Supplemental Figure 4B-F).

Finally, we investigated the impact of GRR presence and position upon L1 ORF1 protein expression levels from natural and synthetic mouse L1 constructs. We transfected HeLa cells with L1spa_Dbl, L1spa_Dbl_SMBB, L1spa_Dbl_SMBB_GRRAN, smL1, and smL1_spa3’UTR and performed Western blots on HeLa cell extracts using an antibody specific to mouse L1 ORF1p (54) and an antibody to nucleolin as a loading control. Among three biological replicate experiments, mean nucleolin-normalized L1spa_Dbl ORF1p levels were ∼2-fold lower compared to L1spa_Dbl_smBB, while L1spa_Dbl_smBB_GRRAN ORF1p levels were increased over L1spa_Dbl_smBB by ∼1.5-fold (Figure 3C-D, Supplemental Figure 6). L1 ORF1p levels from smL1 and smL1_spa3’UTR were elevated ∼15-fold and ∼12-fold, respectively, relative to L1spa_Dbl_smBB, consistent with the previously reported increase in protein expression from synthetic L1 constructs (1, 43). To rule out the absence of a Kozak sequence on the L1spa_Dbl ORF1p initiation codon as the reason for diminished ORF1p levels compared to the other Kozak-containing constructs (Figure 2), we verified in a separate experiment that ORF1p levels from L1spa_Dbl_smBB_Restore_3’UTR, which is identical to L1spa_Dbl_smBB except for the inclusion of the L1spa GRR upstream of the NEO cassette (Figure 2A) also are ∼2-fold lower than L1spa_Dbl_smBB (Supplemental Figure 7). Taken together, our results indicate that placement of the GRR upstream, but not downstream, of the NEO cassette diminishes ORF1p expression, implicating reduced ORF1p levels in the position-dependent effect of the GRR upon natural mouse L1 retrotransposition efficiency.

## Discussion

The highly active smL1 construct and its derivatives have facilitated numerous insights into the mechanism, regulation, and consequences of L1 retrotransposition *in vivo* and *in vitro* (44, 55–65). Here we demonstrate that synthetic reduction in mouse L1 ORF adenosine content is not responsible for the previously reported difference in retrotransposition efficiency of smL1 compared to pTN201/L1spa. We find instead that the elevated activity of smL1 over L1spa arises partly from the nonsynonymous ORF1 substitution H159D (46), and to a larger extent from the presence of the 3’UTR GRR sequence upstream of the NEO cassette in the L1spa construct. Our results resolve a long-standing incongruity regarding the impact of synthetic ORF sequences on L1 retrotransposition efficiency, as the similarly re-coded synthetic human L1 ORFeus_Hs retrotransposes with no more than 3-fold greater efficiency than the native parental element L1_RP_ (43, 66). Thus, while both human and mouse synthetic L1 ORFs yield increased L1 RNA levels relative to native ORF sequences, this elevated expression alone does not effect a proportionate increase in retrotransposition (Figure 1, Figure 2, Supplemental Figure 4)(1, 43). This result is consistent with the modest increase in retrotransposition achieved by augmenting L1 expression with a strong heterologous promoter (19). While regulation of L1 expression is a critical mechanism for limiting endogenous retrotransposition (67), our results suggest that in contexts where L1 expression surpasses a certain threshold, L1 mRNA levels are not the limiting determinant of retrotransposition efficiency. Moreover, our findings emphasize that deleting or reconfiguring portions of the L1 sequence when generating engineered reporter constructs can dramatically impact detected retrotransposition efficiency and complicate interpretation of experimental results.

Our results suggest that the molecular mechanism by which deleting or repositioning the L1spa GRR impacts natural mouse L1 retrotransposition efficiency involves modulation of ORF1 protein levels. Relative to a natural mouse L1 construct lacking the GRR (L1spa_Dbl_smBB), placement of the GRR upstream of the NEO cassette (L1spa_Dbl or L1spa_Dbl_smBB_restore_3’UTR) diminishes ORF1p levels 2-fold, and its placement downstream of the NEO cassette, proximal to the 3’ end of the L1 transcriptional unit (L1spa_Dbl_smBB_GRRAN), increases ORF1p levels ∼1.6-fold (Figure 3C-D, Supplemental Figure 6, Supplemental Figure 7). The position-dependent impact of the GRR on natural mouse L1 ORF1p levels contrasts with our results for L1 RNA levels, which appear to be diminished by the inclusion of the GRR regardless of its placement within the construct (Supplemental Figure 4). ORF1p levels from a synthetic mouse L1 construct lacking the GRR (smL1) are elevated ∼15-fold over the congruent natural L1 construct (L1spa_Dbl_smBB), consistent with enhanced protein expression from codon-optimized ORFs, and are modestly reduced by inclusion of the GRR upstream of the NEO cassette (smL1_spa3’UTR)(Figure 3C-D, Supplemental Figure 6). Critically, all the above-described constructs retrotranspose with similar efficiency except for L1spa_Dbl and L1spa_Dbl_smBB_Restore_3’UTR, which retain only ∼20% relative retrotransposition efficiency (Figure 1-2, Supplemental Figure 3).

We propose that increased ORF1p levels effect a proportionate increase in retrotransposition only up to a certain threshold, which for a natural mouse L1 is exceeded when the GRR is deleted or placed downstream of the NEO cassette. Consistently, due to the dramatically elevated levels of smL1 ORF1p expression achieved via codon optimization, the diminution of ORF1p levels when the GRR is placed upstream of the NEO cassette (smL1_spa3’UTR) has no impact on retrotransposition efficiency (Figure 2A, Figure 3C-D, Supplemental Figure 6). This model also is consistent with the inability of GRR deletion or repositioning to fully overcome the negative effect of the H159 variant in L1spa ORF1p (Figure 2B). While L1spa_smBB and L1spa_smBB_GRRAN likely produce more ORF1p than L1spa_Dbl, the ORF1p produced is impaired for nucleic acid chaperone activity and therefore efficient retrotransposition, rendering the H159 variant largely epistatic to the deleted or repositioned GRR. The mechanism by which the presence and position of the GRR sequence influence L1 ORF1p translation or steady-state levels is a topic for future investigation. In addition, the impact of the GRR on L1 ORF2 protein levels remains to be elucidated.

The GRR sequences from both active and ancient mouse L1 subfamilies diminish natural mouse L1 retrotransposition efficiency when placed upstream of the NEO cassette (Supplemental Figure 3A). Previous reports suggest that the equivalent region of the human L1 3’UTR does not have the same effect on engineered human L1 retrotransposition. The human L1 3’UTR poly-purine tract comprises four runs of 3 to 5 guanine residues across a 34 bp region, which have the demonstrated ability to participate in G-quadruplex secondary structures (22, 68, 69), and in frequently used human L1 reporter constructs this 3’UTR sequence is situated upstream of the NEO cassette (19). In contrast to our results with a natural mouse L1 element, deleting 145 bp of the human 3’UTR including the poly-purine tract in a NEO-based human L1 reporter construct diminished retrotransposition efficiency by ∼10% (19). In a subsequent study, deletion of only the 34 bp poly-purine tract diminished retrotransposition of an EGFP-based L1 reporter construct by ∼30%, as did mutation of 12 guanine residues to ablate G-quadruplex secondary structure formation; small molecule G-quadruplex stabilization also was shown to very modestly stimulate engineered retrotransposition (69). Future studies are required to determine whether human L1 poly-purine tract presence and position within engineered reporter constructs impact L1 RNA expression and protein expression, and how these potential effects are related to the formation and stabilization of G-quadruplex structures.

Taken together, our results suggest that the inhibitory effect of the GRR sequence on retrotransposition is position-dependent and may be specific to engineered L1 reporters. Indeed, the possibility that disruption of the L1 3’UTR by an indicator cassette could impact retrotransposition has long been acknowledged (70) and, to speculate, the conservation of a self-attenuating motif is difficult to reconcile with the evolutionary imperative of L1 as a selfish genetic element (20, 22, 71). Positioning the GRR near the 3’ end of the L1 transcriptional unit, where it is found in endogenous L1 elements, does not elevate retrotransposition efficiency more than deleting the GRR entirely, but it does cause a modest increase in ORF1p expression (Figure 3C-D, Supplemental Figure 6, Supplemental Figure 7). Perhaps this positive influence on ORF1p (and speculatively, perhaps ORF2p) expression is more important for endogenous L1 activity *in vivo* than is reflected in an engineered reporter system. The extent to which the deletion of the GRR from the smL1 construct and its derivatives impacts the conclusions drawn from previous *in vitro* and *in vivo* studies using these reporters remains to be evaluated (44, 55–64); at least some transgenic mouse models generated using smL1 based constructs lack the GRR (56, 60, 63, 64). Engineered mouse L1 constructs reconfigured with the GRR downstream of the reporter cassette represent useful tools for investigating the function of this conserved motif, and for studies of mouse L1 biology in general, as high efficiency retrotransposition and structurally normal L1 integrants can be achieved using a construct containing the entire native mouse L1 sequence.

## Materials and Methods

### Retrotransposition indicator constructs

All constructs used in this study are summarized in Table 1. pTN201/L1spa: Has been described previously (6). It consists of the pCEP4 backbone with the element L1spa truncated at position 7418 in the 3’UTR (removing the native mouse L1 polyadenylation signal but leaving the G-rich region (GRR) of the 3’UTR in place). The L1spa 3’UTR is followed by the mneoI retrotransposition indicator cassette (19, 72). The mneoI cassette is followed by the human L1 polyadenylation signal which is directly upstream of the SV40 polyadenylation signal in the pCEP4 backbone.

**Table 1:** Summary of construct characteristics. From left to right, columns contain the construct name, the type of ORF sequences (natural or synthetic), whether a Kozak sequence is present at the initiation codon of L1 ORF1p, the amino acid at L1 ORF1p position 53, the amino acid at L1 ORF1p position 159, the origin of the 3’UTR non-GRR portion (L1spa or smL1), the origin of the 3’UTR GRR portion (L1spa, other L1 subfamily consensus, or truncated L1spa GRR), and the position of the GRR sequence relative to the NEO cassette.

| Construct Name | ORF sequences | Kozak Sequence? | ORF1- 53 | ORF1-159 | 3'UTR non-GRR | 3'UTR GRR | GRR position |
| --- | --- | --- | --- | --- | --- | --- | --- |
| L1spa | natural | no | G | H | L1spa | L1spa | before NEO |
| L1spa_G53D | natural | no | D | H | L1spa | L1spa | before NEO |
| L1spa_H159D | natural | no | G | D | L1spa | L1spa | before NEO |
| L1spa_Dbl | natural | no | D | D | L1spa | L1spa | before NEO |
| smL1 | synthetic | yes | D | D | smL1 | Deleted | - |
| smL1_D53G | synthetic | yes | G | D | smL1 | Deleted | - |
| smL1_D159H | synthetic | yes | D | H | smL1 | Deleted | - |
| smL1_Dbl | synthetic | yes | G | H | smL1 | Deleted | - |
| L1spa_Dbl_smBB | natural | yes | D | D | smL1 | Deleted | - |
| L1spa_Dbl_smBB_Delete_Kozak | natural | no | D | D | smL1 | Deleted | - |
| L1spa_Dbl_smBB_Restore_3'UTR | natural | yes | D | D | L1spa | L1spa | before NEO |
| smL1_spa3'UTR | synthetic | yes | D | D | L1spa | L1spa | before NEO |
| L1spa_Dbl_smBB_GRRAN | natural | yes | D | D | smL1 | L1spa | after NEO |
| L1spa_smBB | natural | yes | G | H | smL1 | Deleted | - |
| L1spa_smBB_GRRAN | natural | yes | G | H | smL1 | L1spa | after NEO |
| L1spa_Dbl_smBB_Restore_L1spa_nonGRR | natural | yes | D | D | L1spa | Deleted | - |
| L1spa_Dbl_smBB_Restore_L1spa_GRR | natural | yes | D | D | Deleted | L1spa | before NEO |
| L1spa_Dbl_smBB_V_GRR | natural | yes | D | D | Deleted | L1Md_V | before NEO |
| L1spa_Dbl_smBB_F_GRR | natural | yes | D | D | Deleted | L1Md_F | before NEO |
| L1spa_Dbl_smBB_GF_GRR | natural | yes | D | D | Deleted | L1Md_GF | before NEO |
| L1spa_Dbl_smBB_A_GRR | natural | yes | D | D | Deleted | L1Md_A | before NEO |
| L1spa_Dbl_smBB_G1 | natural | yes | D | D | L1spa | Truncated | before NEO |
| L1spa_Dbl_smBB_G2 | natural | yes | D | D | L1spa | Truncated | before NEO |
| L1spa_Dbl_smBB_G3 | natural | yes | D | D | L1spa | Truncated | before NEO |
| L1spa_Dbl_smBB_G4 | natural | yes | D | D | L1spa | Truncated | before NEO |
| L1spa_Dbl_smBB_G5 | natural | yes | D | D | L1spa | Truncated | before NEO |
| L1spa_Dbl_smBB_G6 | natural | yes | D | D | L1spa | Truncated | before NEO |
| L1spa_Dbl_smBB_G7 | natural | yes | D | D | L1spa | Truncated | before NEO |
| L1spa_Dbl_smBB_G8 | natural | yes | D | D | L1spa | Truncated | before NEO |
| L1spa_Dbl_smBB_G9 | natural | yes | D | D | L1spa | Truncated | before NEO |

### L1spa_G53D

Identical to pTN201/L1spa but with glycine 53 of ORF1 replaced by aspartic acid.

### L1spa_H159D

Identical to pTN201/L1spa but with histidine 159 of ORF1 replaced by aspartic acid.

### L1spa_dbl

Identical to pTN201/L1spa but with glycine 53 of ORF1 replaced by aspartic acid and histidine 159 of ORF1 replaced by aspartic acid.

### smL1

Has been described previously (1). It consists of the pCEP4 backbone with a synthetic mouse L1 element based on L1spa. The initiation codon of ORF1 has been placed into an engineered Kozak consensus. The amino acid sequence of smL1 is identical to that of L1spa except for two residues in ORF1 (position 53 and position 159). The 3’UTR of smL1 contains several single nucleotide substitutions relative to L1spa, as well as two single-nucleotide deletions, and is truncated at position 7259 of L1spa. This truncation deletes the GRR of the L1spa 3’UTR. The smL1 3’UTR is followed by the mneoI retrotransposition indicator cassette. The mneoI cassette is followed by the human L1 polyadenylation signal which is directly upstream of the SV40 poly-A signal in the pCEP4 backbone.

### smL1_mut2

Has been described previously (1). It is identical to smL1 but with inactivating mutations in the endonuclease and reverse transcriptase active sites of ORF2.

### smL1_D53G

Identical to smL1 but with aspartic acid 53 of ORF1 replaced by glycine. smL1_D159H: Identical to smL1 but with aspartic acid 159 of ORF1 replaced by histidine.

### smL1_Dbl

Identical to smL1 but with aspartic acid at position 53 of ORF1 replaced by glycine and aspartic acid at position 159 of ORF1 replaced by histidine.

### L1spa_dbl_smBB

“L1spa double; smL1 backbone”. It consists of the L1spa_dbl sequence, with a Kozak consensus at the initiation codon of ORF1 and the mutated 3’UTR of smL1.

L1spa_dbl_smBB _Del_Kozak: Identical to L1spa_dbl_smBB but with the Kozak consensus removed.

L1spa_dbl_smBB_Restore_3’UTR: Identical to L1spa_dbl_smBB but with the mutated 3’UTR of L1SM replaced by the native L1spa 3’UTR as found in pTN201/L1spa.

### L1spa_Dbl_smBB_GRRAN

“L1spa double; smL1 backbone; G-rich region after NEO”. Identical to L1spa_dbl_smBB but with the GRR of L1spa (nucleotides 7260-7416) and an engineered AflII restriction site placed after the mneoI cassette and before the SV40 polyadenylation signal of the pCEP4 backbone, and the human L1 polyadenylation signal deleted.

L1spa_Dbl_SMBB_Restore_L1spaGRR: Identical to L1spa_Dbl_SMBB, but the GRR sequence from the L1spa 3’UTR placed between the end of ORF2 and the NEO cassette, using an existing AgeI site and introducing an engineered SbfI site.

### smL1_L1spa3’UTR

Consists of the smL1 ORFs with a Kozak sequence on smL1 ORF1 and the native L1spa 3’UTR situated upstream of the mneoI cassette.

### L1spa_smBB

Identical to L1spa_Dbl_smBB but with the original L1spa ORF1p amino acid sequence (G53; H159).

### L1spa_smBB_GRRAN

Identical to L1spa_Dbl_smBB_GRRAN except with the original L1spa ORF1p amino acid sequence (G53; H159).

### L1spa_Dbl_smBB_Restore_L1spa_nonGRR

Identical to L1spa_Dbl_smBB, but with the non-GRR portion of the L1spa 3’UTR inserted upstream of the mneoI cassette.

L1spa_Dbl_smBB_V_GRR, L1spa_Dbl_smBB_F_GRR, L1spa_Dbl_smBB_G_F__GRR, and L1spa_Dbl_smBB_A_GRR: Identical to L1spa_Dbl_smBB_L1spaGRR, but with the GRR sequence (but not the non-GRR 3’UTR sequence) from the L1Md_VI, L1Md_FI, L1Md_GFI and L1Md_AI consensus sequences (8) placed between the end of ORF2 and the NEO cassette.

### L1spa_Dbl_SMBB_G1 through L1spa_Dbl_SMBB_G9

Identical to L1spa_Dbl_SMBB, but with increasing portions of the GRR sequence from the L1spa 3’UTR, each restoring an additional run of 3 or more guanine residues, introduced between the end of ORF2 and the NEO cassette as described for L1spa_Dbl_smBB_Restore_L1spa_GRR.

### pCEP4_GFP

Consists of the pCEP4 backbone with a humanized Renilla GFP reporter gene.

### Construct generation

all cloning was performed using restriction enzymes from New England Biolabs (NEB) according to manufacturer’s instructions. PCR fragments were generated using oligonucleotides ordered from Integrated DNA Technologies (IDT) and Q5 High Fidelity DNA Polymerase 2X master mix (NEB), according to manufacturer’s instructions. Constructs were thoroughly capillary sequenced to confirm the absence of random PCR-induced mutations.

### Cell culture

HeLa-JVM cells (19) were maintained in Dulbecco’s Modified Eagle’s Medium (DMEM) high glucose supplemented with 1% L-glutamine, 1% pen-strep and 10% fetal bovine serum. Cells were passaged every 3-5 days at 70-80% confluence using 0.25% Trypsin-EDTA (Life Technologies). Cells were grown at 37°C in 5% CO_2_.

### Retrotransposition assays

For transient retrotransposition assays performed in 6-well plates as previously described (35, 37), 5x10^3^ HeLa-JVM cells were plated per well. 24 hours later, each well was transfected with 1 μg of plasmid DNA using 4 µL FuGene and 96 µL opti-MEM. Each construct was transfected in triplicate. The media was replaced 24 hours post-transfection. At 72 hours post-transfection, selection was initiated with 400 μg/ml G418 (Geneticin, Life Technologies). G418 media was replaced every other day. At 12 days post-transfection, cells were fixed with 2% formaldehyde/0.4% glutaraldehyde solution and stained with 0.1% crystal violet solution.

Assays to measure transfection efficiency for each L1 reporter plasmid were carried out as follows: 4x10^4^ HeLa cells were plated per well of a 6-well dish. 24 hours later, each well was transfected with 1 μg total plasmid DNA, consisting of 0.5 μg L1 reporter plasmid and 0.5 μg pCEP4-GFP, using 4 µL of FuGene and 96 µL opti-MEM. Each L1 reporter+GFP plasmid was transfected in duplicate. The media was replaced 24 hours post-transfection. At 72 hours post-transfection, cells were trypsinized, resuspended in PBS (Life Technologies) and assessed for GFP expression by flow cytometry using a Cytoflex flow cytometer (Translational Research Institute flow cytometry core). The average percent of GFP-positive cells was determined for each L1 reporter construct, and used to normalize the results of the retrotransposition assay.

### qRT-PCR

2x10^5^ HeLa cells were plated per well of a 6-well dish and transfected 24 hours later with 1 µg of L1 reporter plasmid using 4 µL of FuGene and 96 µL of opti-MEM. Media was replaced 24 hours post-transfection with media containing 200 µg/mL Hygromycin for 8 days total selection. RNA was extracted using the RNAeasy mini kit (Qiagen) from HeLa cells at 2 days post transfection. Independently transfected wells were considered as biological replicates. RNA was treated with the Turbo DNA free kit (Invitrogen) using the rigorous protocol. cDNA was generated with the iScript cDNA synthesis kit using 1 µg of input RNA. qPCR was carried out using PowerUp SYBR Green Master Mix (A25742 Thermo Fisher) We calculated the average Ct value across three technical replicates for each biological replicate. The delta Ct value (averageCt gene - averageCt RPS18) was used to determine the expression relative to RPS18 whilst accounting for the amplification efficiency of each primer set (efficiency value^-ΔCt^). Primer efficiencies were determined using serial cDNA dilutions to generate a non-linear regression of the average Ct values for three replicates for each dilution. The slope was used to calculate primer efficiency using the following equation (10(− 1/𝑠𝑙𝑜𝑝𝑒)−1)×100.

### 3’RACE

For each construct, 1x10^5^ HeLa-JVM cells were plated per well of a 6-well dish and transfected ∼18 hours later with 1 µg of plasmid DNA using 4 µL of FugeneHD and 96 µL of Opti-MEM. Cells were selected for 7 days using 200 µg/ml of Hygromycin B, and total RNA was extracted using the Qiagen RNeasy Mini kit (cat no. 74104). 3’RACE was performed using the 3’RACE module of the Invitrogen First Choice RLM-RACE kit (cat no. AM1700) according to manufacturer’s instructions. Briefly, 1ug of total RNA was used in a reverse transcription reaction with kit provided M-MLV reverse transcriptase, RNase inhibitor, and 3’RACE adapter (5’-GCGAGCACAGAATTAATACGACTCACTATAGGT12VN-3’). RNA was reverse transcribed at 42°C for 1 hour. PCR amplification of 3’RACE products was performed using gene-specific primers to the 5’ region of the L1spa 3’UTR (3’RACE_3’UTR-3 CGAAAGGACCCAGATGTAGC), the antisense SV40 early promoter within the NEO retrotransposition indicator cassette (asSV40prom_F CGGGACTATGGTTGCTGACT), and the 3’ region of the hygromycin resistance gene in the pCEP4 plasmid backbone (Hygro_F3 ATTTCGGCTCCAACAATGTC) in combination with the adaptor specific primer from the RLM-RACE kit (3’RACE Outer Primer 5’-GCGAGCACAGAATTAATACGACT-3’). PCR cycling conditions were as follows: 3 minutes at 94°C, 35 cycles of 94°C 30 seconds, 55°C 30 seconds, 72°C 30 seconds followed by 7 minutes at 72°C. PCR products were run on a 2% agarose gel, and bands were excised and purified using the Qiagen MinElute gel extraction kit (cat no. 28604). Purified products were cloned using TA Cloning Kit (Invitrogen, K204001.) and sequenced using primers T7 TAATACGACTCACTATAGGG and Neosplice_F2 AGTGACAACGTCGAGCACAG by AGRF Brisbane.

### L1 insertion mapping and characterization

Clonal cell lines for inverse PCR characterization of L1spa_Dbl_smBB_GRRAN insertions were generated by plating 5x10^3^ HeLa cells per 10 cm dish, and 18 hours later transfecting with 1 µg plasmid DNA using 4 µL FuGene and 96 µL opti-MEM. Media was replaced 24 hours after transfection. Selection with 400 µg/ml Geneticin was initiated at 72 hours post-transfection and continued for 12 days to generate isolated colonies harboring retrotransposition events. Individual colonies were manually picked and expanded to generate clonal cell lines. Approximately 5x10^6^ cells were harvested per clonal cell line and subjected to phenol-chloroform DNA extraction. DNA concentration was measured via NanoDrop. Inverse PCR was carried out as described previously (73, 74). Briefly, 4 µg from each cell line was digested with 25 units EcoRI (New England Biolabs) in 100 µL total volume. Reactions were heat inactivated and digested DNA was ligated under dilute conditions to promote intramolecular ligation (1 ml total volume) using 3200 units T4 DNA ligase (NEB) for 24 hours. Ligations were purified by chloroform extraction and DNA was resuspended in 40 µL water. For each ligation, 4 µL of DNA was used as a template in the first-round inverse PCR reaction using the Roche Expand long-template PCR system. Each reaction consisted of 10 pmol primer NEO210as (5’ GACCGCTTCCTCGTGCTTTACG 3’), 10 pmol primer NEO1720s (5’ TGCGCTGACAGCCGGAACACG 3’), 20 nM each dNTP, and 2.5 units of enzyme. Cycling conditions were 95°C for 2 minutes, followed by 30 cycles of 94°C for 15 seconds, 64°C for 30s, and 68°C for 15 min, followed by a 30 min extension at 68°C. For the second-round inverse PCR reaction, 4 µL of the product from the first-round inverse PCR was used directly as template, with 10 pmol primer NEO173as (5’ CATCGCCTTCTATCGCCTTCTTG 3’) 10 pmol primer NEO1808s (5’ GCGTGCAATCCATCTTGTTCAATG 3’), 20 nM each dNTP, and 2.5 units of enzyme. Cycling conditions were identical to first-round inverse PCR. Products were run on a 1% agarose gel and bands were excised and purified using the Qiagen Min-Elute gel extraction kit. Bands <2kb were cloned using the pGEM-T Easy kit (Promega) and bands >2kb were cloned using the Topo-XL2 PCR kit (Life Technologies) following manufacturer’s instructions. Cloned PCR products were sequenced using M13 forward and reverse primers Australian Genome Research Facility, Brisbane). Validation primers (Integrated DNA Technologies) were designed based on the putative genomic location of each insertion detected by inverse PCR, and insertions were validated as empty/filled and/or by 5’ and 3’ junction PCRs using MyTaq HS DNA polymerase (Bioline). Validation products were run on 1% agarose gels, purified using the Qiagen Min-Elute gel extraction kit, and capillary sequenced (Australian Genome Research Facility, Brisbane) to verify the genomic location and structural hallmarks of each insertion.

### Western blots

HeLa-JVM cells were plated at 2 x 10^5^ cells per well of a 6 well plate. The following day cells were transfected with 1 µg of plasmid DNA using 4 µL of FuGene and 96 µL of opti-MEM. Independently transfected wells were considered as biological replicates. Media was replaced within 24 hours of transfection with media containing Hygromycin B (Invitrogen 10687-010), 200 µg/mL. Following 8 days of Hygromycin B selection, cells from each well were collected and lysed using RIPA buffer (Thermo 89900) with Halt protease inhibitor (Thermo 78420). Protein concentration was quantified using the Qubit Protein Assay Kit. 25 µg of sample was diluted in 4x Laemmli buffer with β-mercaptoethanol (Biorad) and heated at 95 degrees Celsius for 5 minutes. Samples were separated on a mini protean gel (4 – 20% Biorad 4561094) and transferred to a PVDF membrane using the iBlot2 system (Invitrogen). Membranes were blocked for 2 hours in Odyssey block (Li-Cor) and incubated with the following antibodies overnight at 4 degrees: anti-ORF1p 1:1000 (EPR21844-108 Abcam), anti-nucleolin 1:2000 (Thermo MA544735). Membranes were washed in PBS + 0.15 Tween20 (PBST) and incubated with IRDye goat anti rabbit 800 1:2000 (Li-Cor 926-68071) for 2 hours at room temperature. Signals were detected using the Odyssey CLx imaging system (Li-Cor). Signal density for each band was determined in Fiji and normalized to the nucleolin signal.

## Acknowledgements

HeLa-JVM cells were a gift from John V. Moran. The pTN201/L1spa construct was a gift from Haig H. Kazazian. We thank members of the Richardson and Faulkner laboratories for helpful discussion and critical reading of the manuscript. J.D.B. is a Founder and Director of CDI Labs, Inc., a Founder of Neochromosome, Inc, a Founder and SAB member of ReOpen Diagnostics, and serves or served on the Scientific Advisory Board of the following: Sangamo, Inc., Modern Meadow, Inc., Sample6, Inc. and the Wyss Institute.

---

**Supplemental Figure 1:**
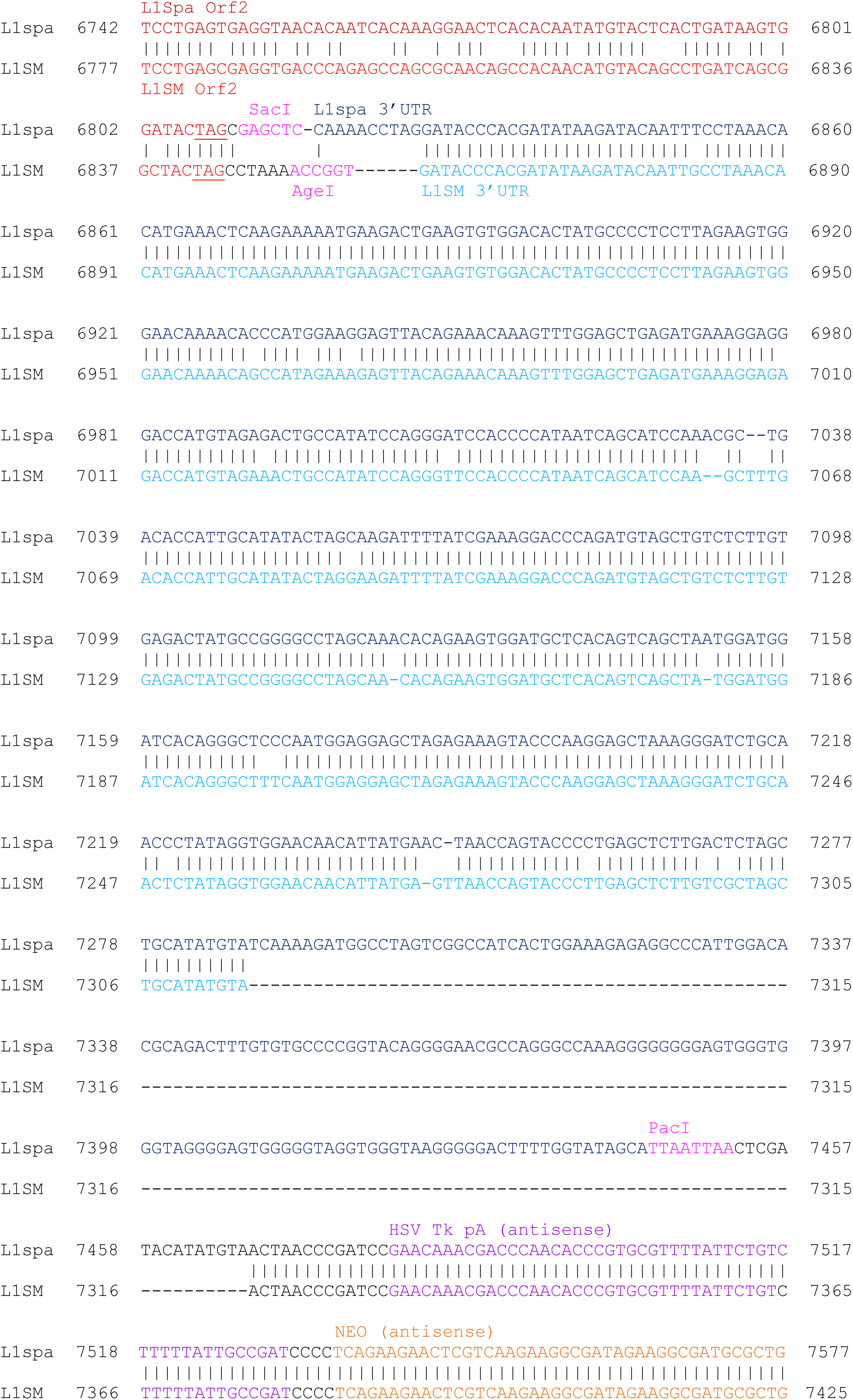
Alignment of the 3’UTRs and flanking sequence of pTN201/L1spa and smL1. Alignment depicts the 3’ end of ORF2 (L1spa ORF2, dark red; smL1 ORF2, bright red), followed by the 3’UTR (L1spa 3’UTR dark blue; smL1 3’UTR, light blue). The smL1 3’UTR deletion is highlighted in yellow, and the runs of guanine bases in the L1spa 3’UTR are highlighted in red. The positions of engineered restriction sites in each construct are shown in pink, and the NdeI sites resulting in the GRR deletion are shown in green. The HSV Tk polyadenylation signal belonging to the NEO cassette (purple) and the second exon of the NEO cassette (orange) are identical between pTN201/L1spa and smL1.

**Supplemental Figure 2:**
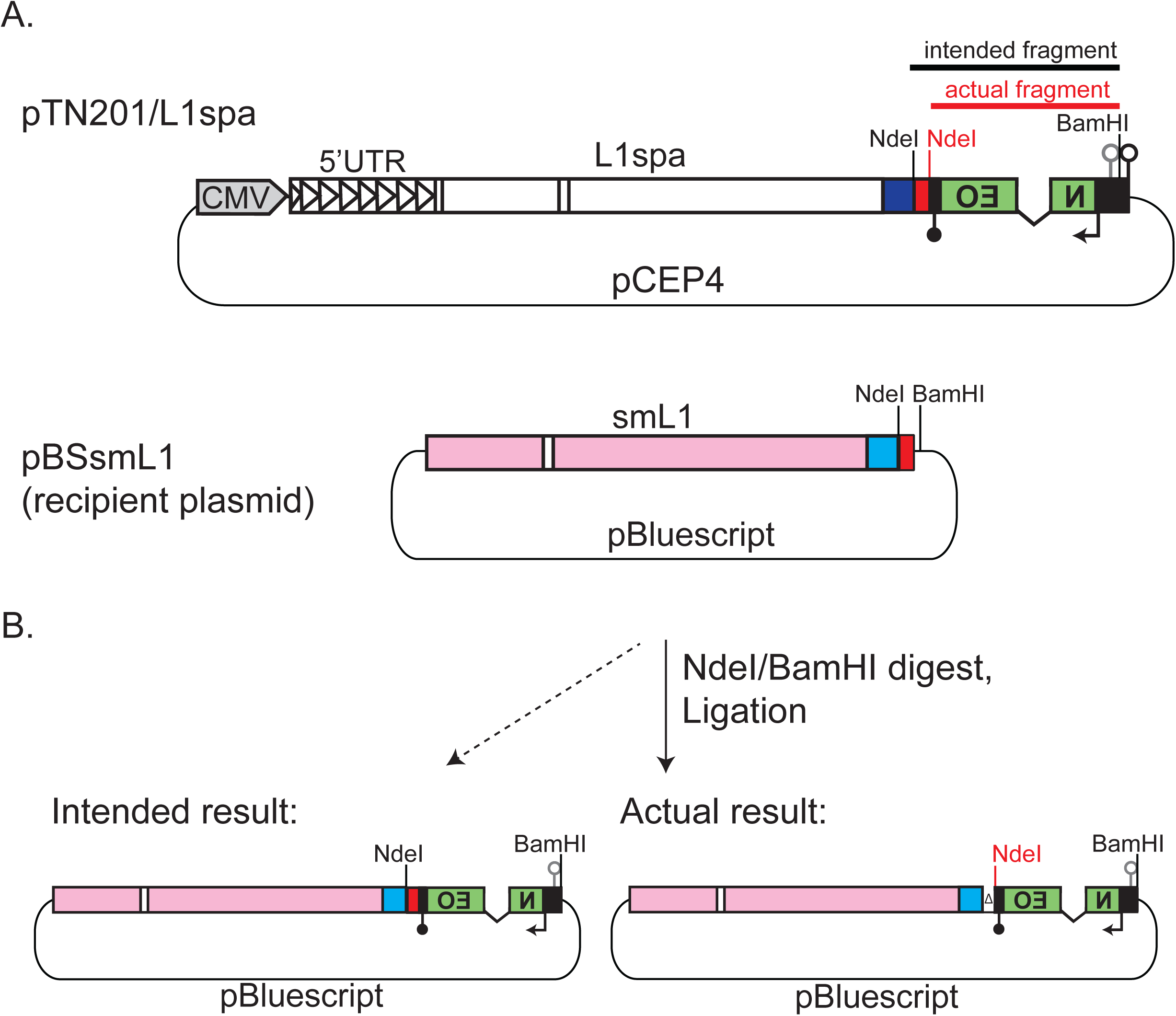
Deletion of the GRR during cloning of the smL1 reporter construct. A. The pTN201/L1spa construct (6) and an intermediate pBluescript clone containing the smL1 sequence, with features annotated as in Figure 1. The positions of known NdeI and BamHI sites are shown in black. Our map of pTN201 did not include 15 base pairs of human L1 sequence present in pTN201 immediately following the L1spa 3’UTR, which was carried over from subcloning described in (6). This 15 bp of human L1 sequence contained an NdeI site (shown in red). B) Deletion of the GRR-containing NdeI fragment during cloning. Left: we intended to move an NdeI/BamHI fragment containing the GRR of the L1spa element and the NEO cassette downstream of the smL1 ORFs and 5’ portion of the 3’UTR in pBluescript. Right: due to the unanticipated NdeI site, a 180 bp NdeI fragment containing the terminal GRR-containing 159 bp of the L1spa 3’UTR and an engineered PacI site (not shown) was deleted from the resulting construct. Owing to its small size, the deletion was not detected when screening the constructs with diagnostic restriction digests.

**Supplemental Figure 3:**
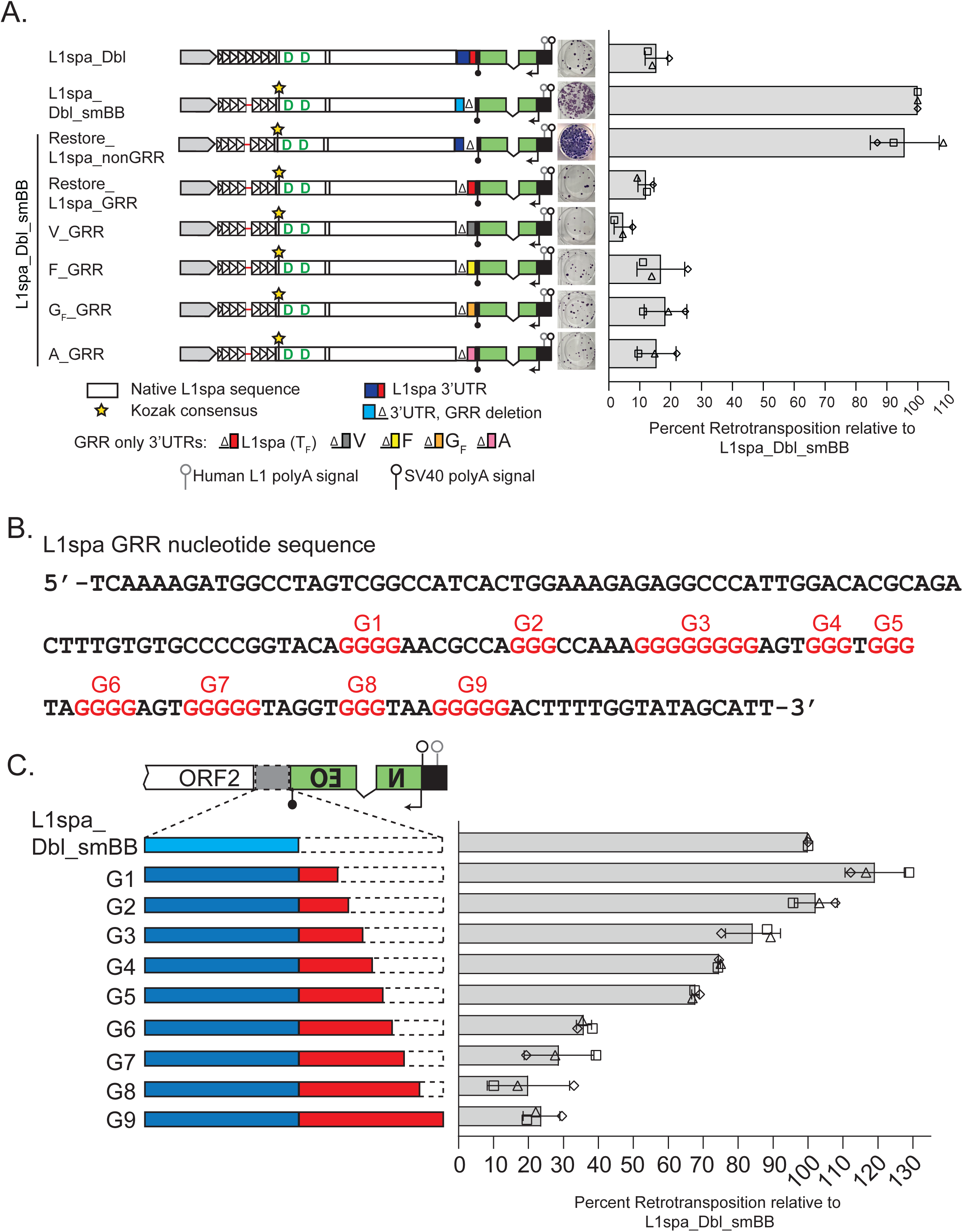
Variations in G-rich region sequence content differentially impact retrotransposition. A. From left to right: a schematic of each construct, a representative well from the retrotransposition assay, and its retrotransposition efficiency relative to smL1. L1spa_Dbl_smBB_Restore_L1spa_nonGRR contains the non-GRR portion only of the L1spa 3’UTR. L1spa_Dbl_smBB_Restore_L1spa_GRR contains the GRR portion only of the L1spa 3’UTR. L1Spa_Dbl_smBB_V_GRR, F_GRR, GF_GRR and A_GRR contain the GRR portion of the 3’UTR from the L1Md_V_I_, L1Md_F_I_, L1Md_G_FI_ and L1Md_A_I_ consensus sequences. The histogram displays the mean of three biological replicate assays, each comprising three technical replicates per construct. Triangle, square, and diamond shapes indicate the individual values for each biological replicate, and the error bars represent standard deviation among biological replicates. B. Nucleotide sequence of the 159bp region comprising the G-rich region (GRR) of the L1spa 3’UTR. Runs of 3 or more consecutive G nucleotides are denoted as G1 through G9 and highlighted in red. C. Retrotransposition assay of L1spa_Dbl_smBB 3’UTR deletion variants. Runs of guanine residues have been progressively restored to the GRR of constructs G1 through G8, with the full GRR present in construct G9. The histogram displays the mean of three biological replicate assays, each comprising three technical replicates per construct. Triangle, square, and diamond shapes indicate the individual values for each biological replicate, and the error bars represent standard deviation among biological replicates.

**Supplemental Figure 4:**
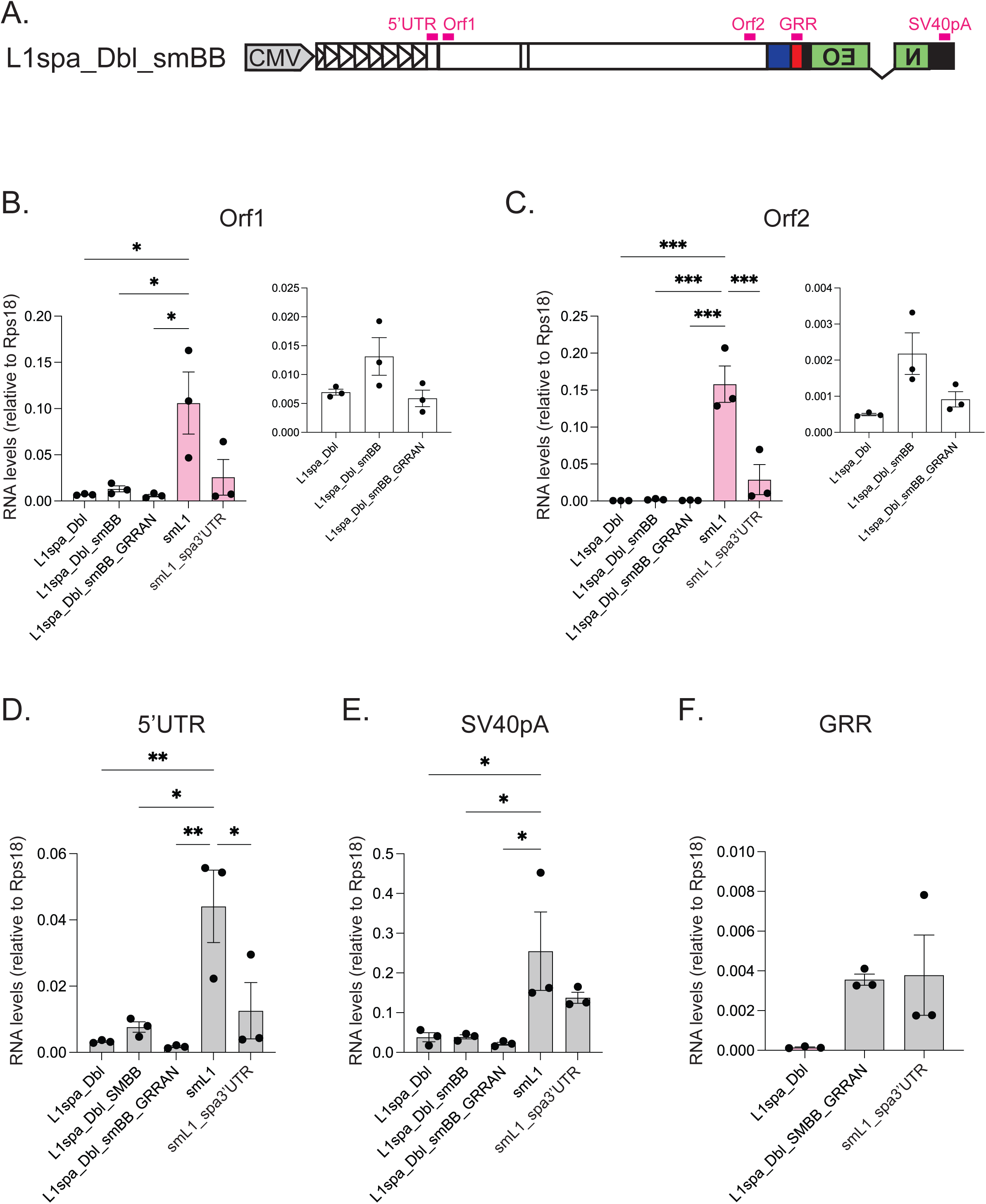
Synthetic ORF sequences and the presence and placement of the L1spa GRR influence RNA expression levels from L1 reporter constructs. A) Illustration of the position of qRT-PCR primer pairs, shown along the L1spa_Dbl_smBB construct. The GRR targeting primers do not anneal within L1spa_Dbl_smBB or smL1_spa3’UTR and in L1spa_Dbl_smBB_GRRAN, the GRR targeting primers anneal downstream of the NEO cassette but upstream of the SV40 polyadenylation signal. For smL1 and smL1_spa3’UTR, primers specific to the synthetic smL1 sequence were used for Orf1 and Orf2. B) qRT-PCR quantitation of L1 Orf1 RNA levels relative to the Rps18 housekeeping gene. Constructs assayed are shown on the x-axis and RNA levels relative to Rps18 are shown on the y-axis. Primers annealing to the natural mouse L1 Orf1 sequence were used for L1spa_Dbl, L1spa_Dbl_smBB, and L1spa_Dbl_smBB_GRRAN (white bars); primers annealing to the synthetic mouse L1 Orf1 sequence were used for smL1 and smL1_spa3’UTR (pink bars). Each dot represents a biological replicate (cDNA template generated from an independently transfected well) consisting of the mean of three technical replicates (independent qRT-PCR reactions performed with the same cDNA template). At right, the inset displays the data for natural mouse L1 constructs with the y-axis scaled for improved visibility. Error bars indicate the standard deviation among biological replicates. One-way ANOVA with Tukey’s multiple comparisons test; p<0.05*; p<0.01*; p<0.001*. C) qRT-PCR quantitation of L1 Orf2 RNA levels relative to the Rps18 housekeeping gene, as described in B); primers annealing to the natural mouse L1 Orf2 sequence were used for L1spa_Dbl, L1spa_Dbl_smBB, and L1spa_Dbl_smBB_GRRAN (white bars); primers annealing to the synthetic mouse L1 Orf2 sequence were used for smL1 and smL1_spa3’UTR (pink bars). D) qRT-PCR quantitation of L1 5’UTR RNA levels relative to the Rps18 housekeeping gene. As described in B) except the same primer set was used for all constructs. E) qRT-PCR quantitation of SV40pA RNA levels relative to the Rps18 housekeeping gene, as described in D. F) qRT-PCR quantitation of L1spa GRR RNA levels relative to the Rps18 housekeeping gene, as described in D. The GRR sequence is not present in the L1spa_Dbl_smBB and smL1 constructs, and for these constructs no qPCR signal was detected for this target.

**Supplemental Figure 5:**
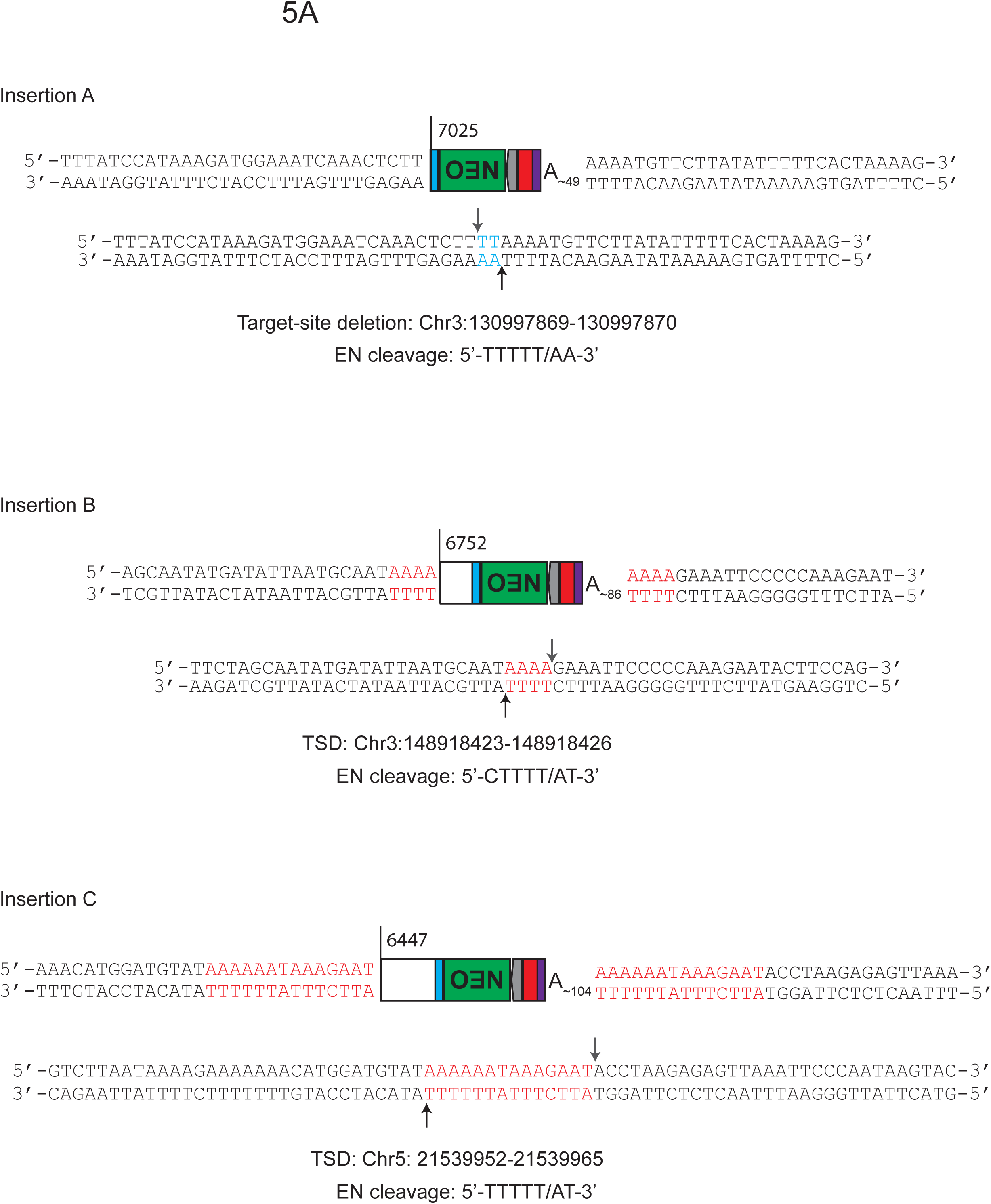
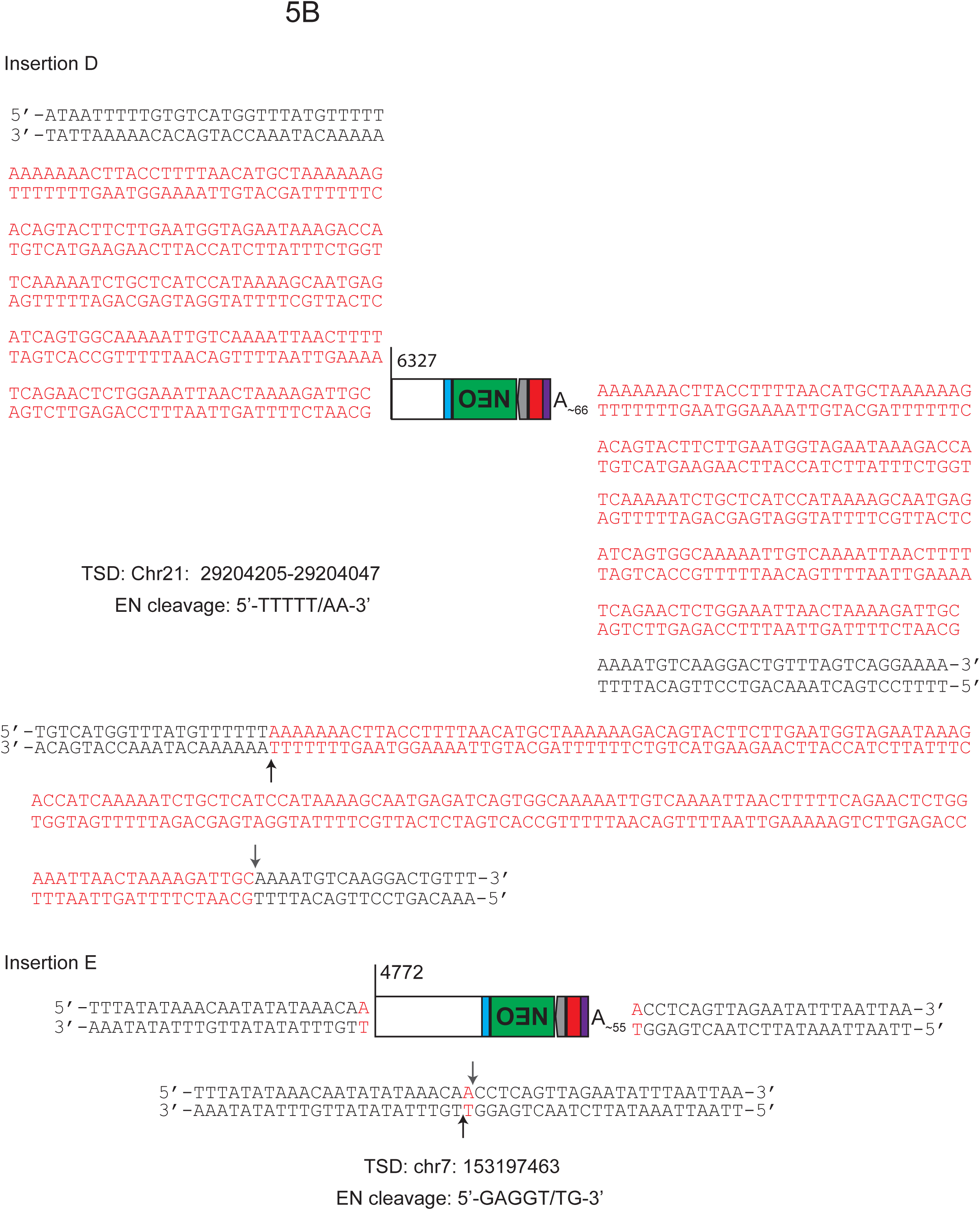
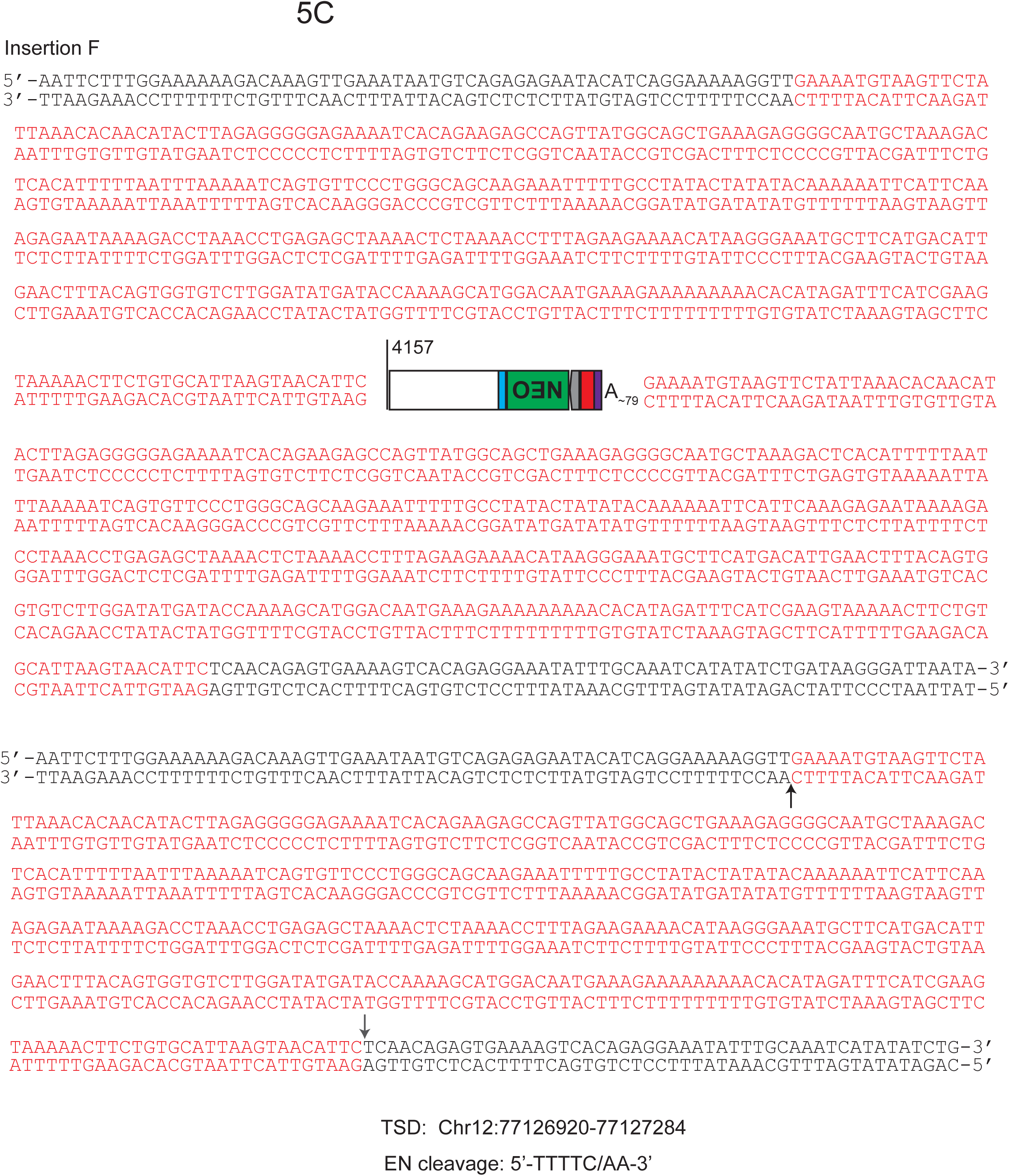
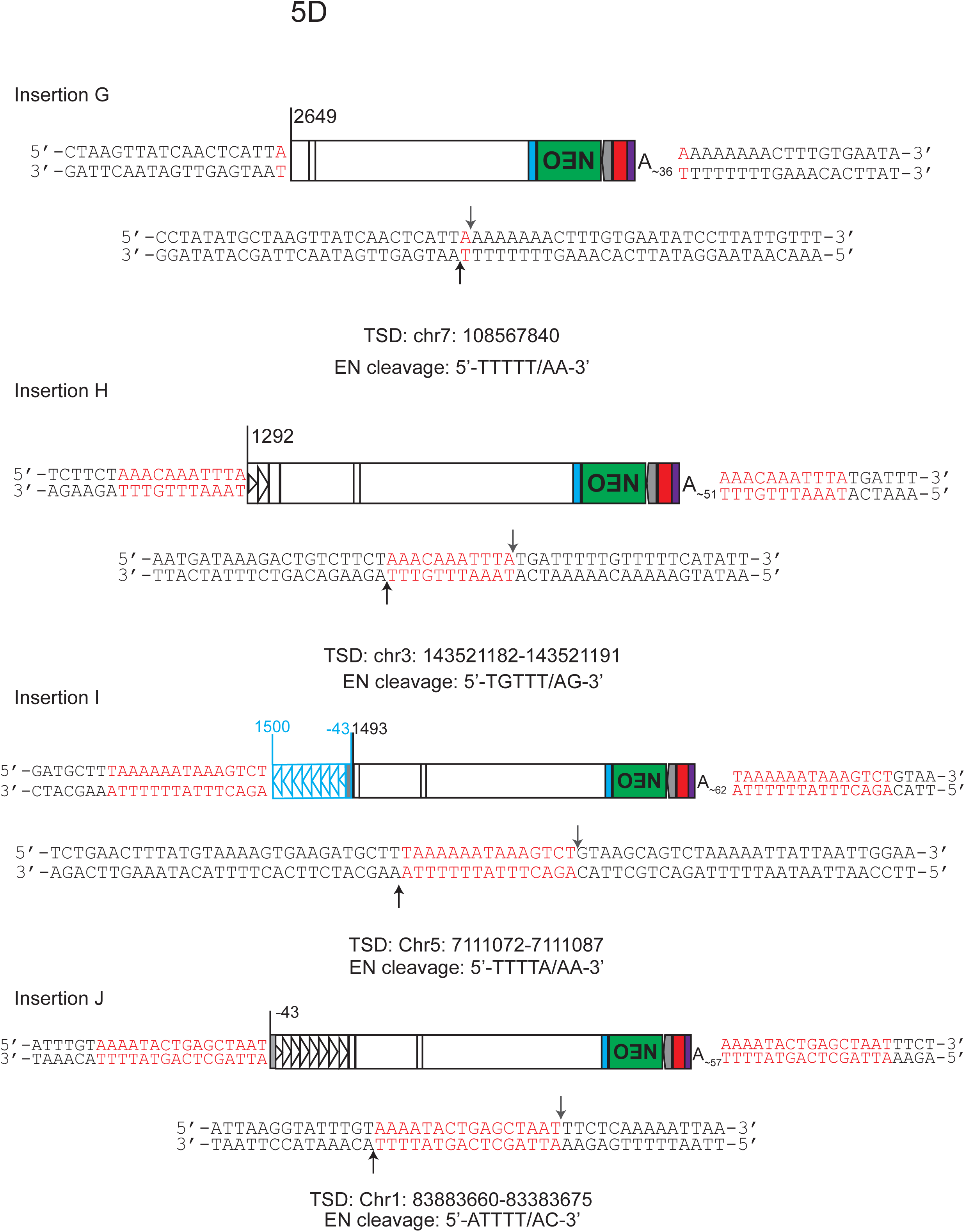
Detailed structures of L1spa_Dbl_smBB_GRRAN integrants characterized via inverse PCR. The structure of each insertion is presented as follows: L1spa 5’UTR (triangles), ORFs (white rectangles), first part of the smL1 3’UTR (light blue), spliced NEO cassette (green), sv40 promoter (grey), GRR (red) sv40 polyadenylation signal (purple). The 5’ truncation point for each insertion is shown. Insertions I and J arose from a transcript initiating upstream of the L1 5’UTR, at the transcription start site of the CMV promoter within the pCEP4 vector (75) denoted as position -43. Insertion J had a 5’ inversion with breakpoint at position 1493 shown. Flanking genomic DNA is shown to the right and left, with target-site duplications in red. The empty site is shown below, with TSD sequence in red, the first-strand cleavage site indicated by a black arrow and the second-strand cleavage site indicated by a grey arrow. Insertion A had a 2 bp target-site deletion, depicted in blue. The chromosomal coordinates of the TSD sequence and the endonuclease cleavage motif for each insertion is shown.

**Supplemental Figure 6:**
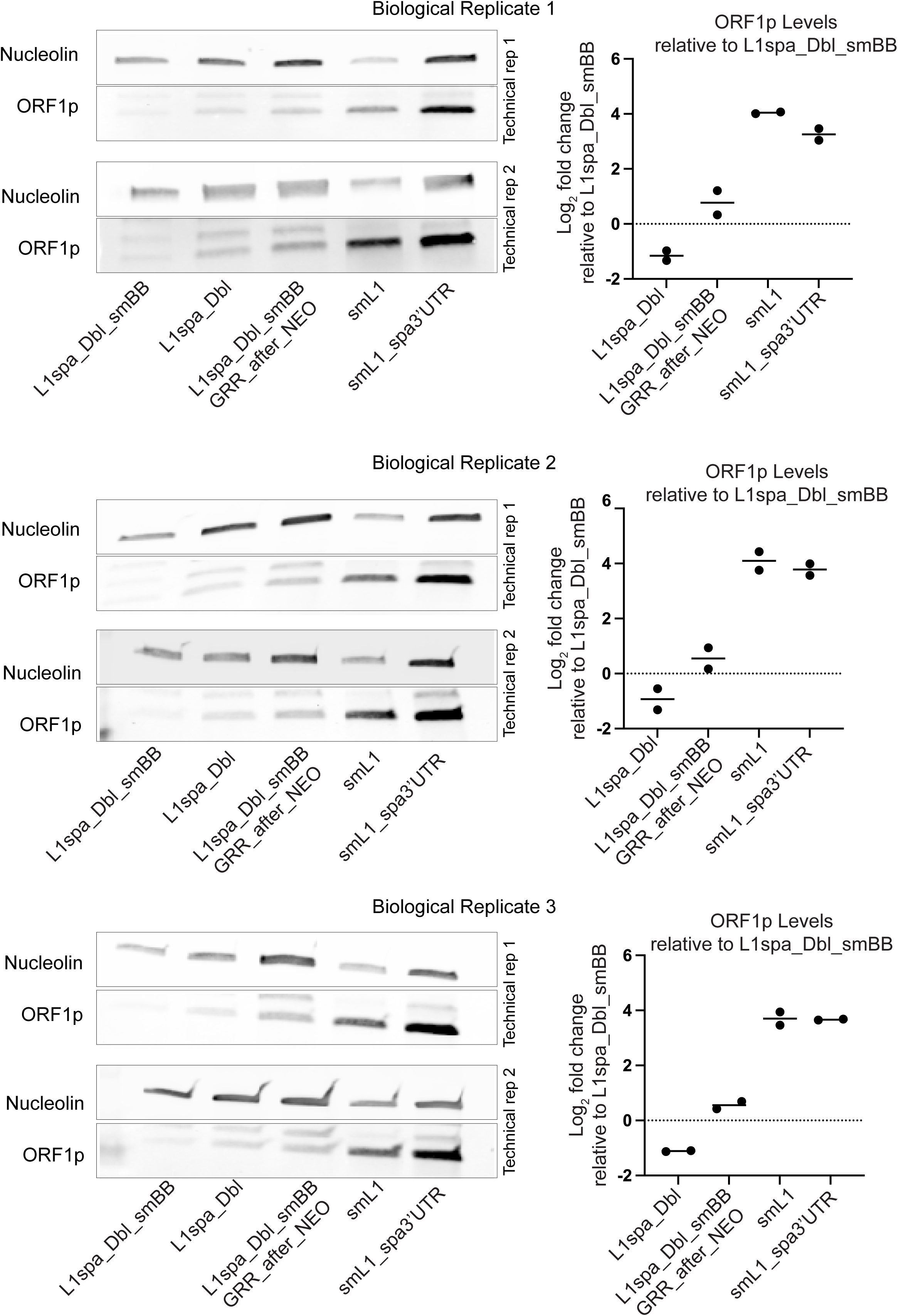
Individual Western blots used in the quantitation of L1 ORF1p levels. For each construct, each biological replicate constitutes an independent transfection. For each biological replicate, two independent blots were performed as technical replicates. For each biological replicate, gel images of the corresponding technical replicates are shown at left. At right, for each biological replicate, Log_2_ fold change in nucleolin-normalized band intensity relative to L1spa_Dbl_SMBB is plotted for L1spa_Dbl, L1spa_Dbl_smBB_GRRAN, smL1, and smL1_spa3’UTR. Each dot represents a technical replicate; horizontal line represents the mean of technical replicates and error bars represent SEM among technical replicates. Technical Replicate 1 from Biological Replicate 1 is used as the representative blot in Figure 3D.

**Supplemental Figure 7:**
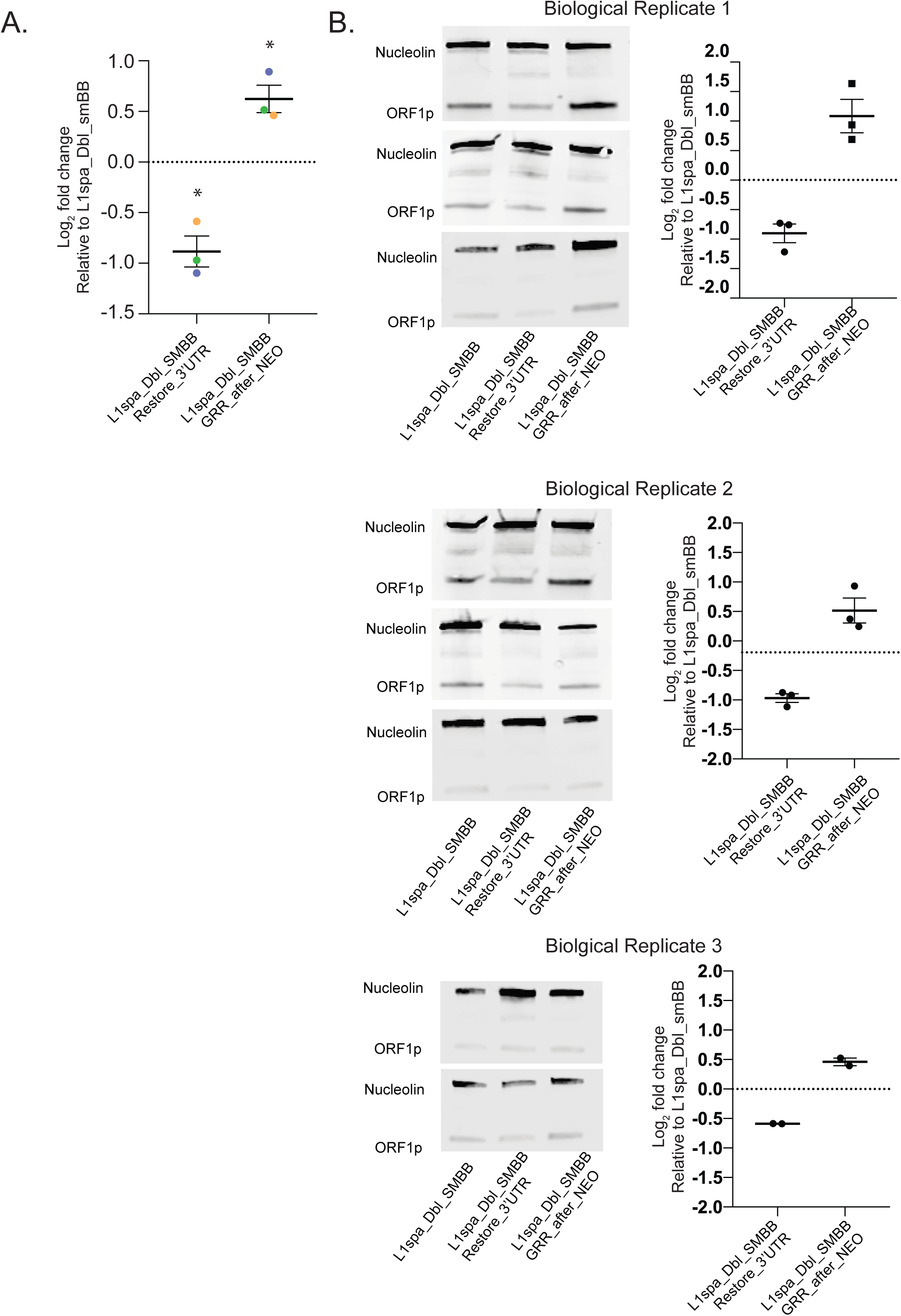
Individual Western blots for quantitation of L1spa_Dbl_smBB_restore_3’UTR ORF1p levels. A. Quantification of nucleolin-normalized ORF1p expression as Log_2_ fold change for L1spa_Dbl_smBB_restore_3’UTR and L1spa_Dbl_SMBB_GRR_after_NEO relative to L1spa_Dbl_smBB. Each biological replicate (cell lysate from an independent transfection) is represented by a colored dot (rep 1=blue, rep 2=green, rep 3=yellow). Horizontal line shows the mean of biological replicates. Values for each biological replicate are the mean of two or three technical replicates. One sample t and Wilcoxon test; p<0.05*. B. Individual Western blots used in the quantitation of L1 ORF1p levels shown in A. For each construct, each biological replicate constitutes an independent transfection. For each biological replicate, two or three independent blots were performed, each blot constituting a technical replicate. For each biological replicate, gel images of the corresponding technical replicates are shown at left. At right, for each biological replicate, Log_2_ fold change in nucleolin-normalized band intensity relative to L1spa_Dbl_SMBB is plotted for L1spa_Dbl_smBB_restore_3’UTR and L1spa_Dbl_smBB_GRRAN. Each dot represents a technical replicate; horizontal line represents the mean of technical replicates and error bars represent SEM among technical replicates.

**Supplemental Table 1:** The first tab contains the sequences of all constructs used in this study, determined by Oxford Nanopore Technologies long-read sequencing. The second tab contains the sequences (iPCR_L1_rev and iPCR_L1_fwd) of inverse PCR products characterizing of L1spa_Dbl_smBB_GRRAN integrants and the corresponding validation primers. The third tab contains retrotransposition assay data (average colony counts from three technical replicate wells, normalized to transfection efficiency) and mean/standard deviation values for three biological replicates used to generate the graphs shown in Figure 1C, Figure 2, and Figure 4 A and C.

## References

1. Han JS, Boeke JD. A highly active synthetic mammalian retrotransposon. Nature. 2004;429(6989):314–8.

2. Kazazian HH, Jr., Moran JV. Mobile DNA in Health and Disease. N Engl J Med. 2017;377(4):361–70.

3. Richardson SR, Doucet AJ, Kopera HC, Moldovan JB, Garcia-Perez JL, Moran JV. The influence of LINE-1 and SINE retrotransposons on mammalian genomes. Microbiology Spectrum. 2015;3(2).

4. DeBerardinis RJ, Goodier JL, Ostertag EM, Kazazian HH, Jr. Rapid amplification of a retrotransposon subfamily is evolving the mouse genome. Nat Genet. 1998;20(3):288–90.

5. Goodier JL, Ostertag EM, Du K, Kazazian HH, Jr. A novel active L1 retrotransposon subfamily in the mouse. Genome Res. 2001;11(10):1677–85.

6. Naas TP, DeBerardinis RJ, Moran JV, Ostertag EM, Kingsmore SF, Seldin MF, et al. An actively retrotransposing, novel subfamily of mouse L1 elements. EMBO J. 1998;17(2):590–7.

7. Penzkofer T, Jager M, Figlerowicz M, Badge R, Mundlos S, Robinson PN, et al. L1Base 2: more retrotransposition-active LINE-1s, more mammalian genomes. Nucleic Acids Res. 2017;45(D1):D68–D73.

8. Sookdeo A, Hepp CM, McClure MA, Boissinot S. Revisiting the evolution of mouse LINE-1 in the genomic era. Mobile DNA. 2013;4(1):3.

9. Zhou M, Smith AD. Subtype classification and functional annotation of L1Md retrotransposon promoters. Mobile DNA. 2019;10:14.

10. Feng Q, Moran JV, Kazazian HH, Jr., Boeke JD. Human L1 retrotransposon encodes a conserved endonuclease required for retrotransposition. Cell. 1996;87(5):905–16.

11. Khazina E, Truffault V, Buttner R, Schmidt S, Coles M, Weichenrieder O. Trimeric structure and flexibility of the L1ORF1 protein in human L1 retrotransposition. Nat Struct Mol Biol. 2011;18(9):1006–14.

12. Khazina E, Weichenrieder O. Non-LTR retrotransposons encode noncanonical RRM domains in their first open reading frame. Proc Natl Acad Sci U S A. 2009;106(3):731–6.

13. Kolosha VO, Martin SL. In vitro properties of the first ORF protein from mouse LINE-1 support its role in ribonucleoprotein particle formation during retrotransposition. Proc Natl Acad Sci U S A. 1997;94(19):10155–60.

14. Kolosha VO, Martin SL. High-affinity, non-sequence-specific RNA binding by the open reading frame 1 (ORF1) protein from long interspersed nuclear element 1 (LINE-1). J Biol Chem. 2003;278(10):8112–7.

15. Luan DD, Korman MH, Jakubczak JL, Eickbush TH. Reverse transcription of R2Bm RNA is primed by a nick at the chromosomal target site: a mechanism for non-LTR retrotransposition. Cell. 1993;72(4):595–605.

16. Mathias SL, Scott AF, Kazazian HH, Jr., Boeke JD, Gabriel A. Reverse transcriptase encoded by a human transposable element. Science. 1991;254(5039):1808–10.

17. Baldwin ET, van Eeuwen T, Hoyos D, Zalevsky A, Tchesnokov EP, Sanchez R, et al. Structures, functions and adaptations of the human LINE-1 ORF2 protein. Nature. 2024;626(7997):194–206.

18. Thawani A, Ariza AJF, Nogales E, Collins K. Template and target-site recognition by human LINE-1 in retrotransposition. Nature. 2024;626(7997):186–93.

19. Moran JV, Holmes SE, Naas TP, DeBerardinis RJ, Boeke JD, Kazazian HH, Jr. High frequency retrotransposition in cultured mammalian cells. Cell. 1996;87(5):917–27.

20. Boissinot S, Sookdeo A. The Evolution of LINE-1 in Vertebrates. Genome Biol Evol. 2016;8(12):3485–507.

21. Goodier JL, Ostertag EM, Kazazian HH, Jr. Transduction of 3’-flanking sequences is common in L1 retrotransposition. Hum Mol Genet. 2000;9(4):653–7.

22. Howell R, Usdin K. The ability to form intrastrand tetraplexes is an evolutionarily conserved feature of the 3’ end of L1 retrotransposons. Mol Biol Evol. 1997;14(2):144–55.

23. Moran JV, DeBerardinis RJ, Kazazian HH, Jr. Exon shuffling by L1 retrotransposition. Science. 1999;283(5407):1530–4.

24. Pickeral OK, Makalowski W, Boguski MS, Boeke JD. Frequent human genomic DNA transduction driven by LINE-1 retrotransposition. Genome Res. 2000;10(4):411–5.

25. Cost GJ, Boeke JD. Targeting of human retrotransposon integration is directed by the specificity of the L1 endonuclease for regions of unusual DNA structure. Biochemistry. 1998;37(51):18081–93.

26. Gilbert N, Lutz S, Morrish TA, Moran JV. Multiple fates of L1 retrotransposition intermediates in cultured human cells. Mol Cell Biol. 2005;25(17):7780–95.

27. Gilbert N, Lutz-Prigge S, Moran JV. Genomic deletions created upon LINE-1 retrotransposition. Cell. 2002;110(3):315–25.

28. Jurka J. Sequence patterns indicate an enzymatic involvement in integration of mammalian retroposons. Proc Natl Acad Sci U S A. 1997;94(5):1872–7.

29. Sultana T, van Essen D, Siol O, Bailly-Bechet M, Philippe C, Zine El Aabidine A, et al. The Landscape of L1 Retrotransposons in the Human Genome Is Shaped by Pre-insertion Sequence Biases and Post-insertion Selection. Mol Cell. 2019;74(3):555-70 e7.

30. Flasch DA, Macia A, Sanchez L, Ljungman M, Heras SR, Garcia-Perez JL, et al. Genome-wide de novo L1 Retrotransposition Connects Endonuclease Activity with Replication. Cell. 2019;177(4):837–51 e28.

31. Feusier J, Watkins WS, Thomas J, Farrell A, Witherspoon DJ, Baird L, et al. Pedigree-based estimation of human mobile element retrotransposition rates. Genome Res. 2019;29(10):1567–77.

32. Richardson SR, Gerdes P, Gerhardt DJ, Sanchez-Luque FJ, Bodea GO, Munoz-Lopez M, et al. Heritable L1 retrotransposition in the mouse primordial germline and early embryo. Genome Res. 2017;27(8):1395–405.

33. Gagnier L, Belancio VP, Mager DL. Mouse germ line mutations due to retrotransposon insertions. Mobile DNA. 2019;10:15.

34. Hancks DC, Kazazian HH, Jr. Roles for retrotransposon insertions in human disease. Mobile DNA. 2016;7:9.

35. Wei W, Morrish TA, Alisch RS, Moran JV. A transient assay reveals that cultured human cells can accommodate multiple LINE-1 retrotransposition events. Anal Biochem. 2000;284(2):435–8.

36. Ostertag EM, Prak ET, DeBerardinis RJ, Moran JV, Kazazian HH, Jr. Determination of L1 retrotransposition kinetics in cultured cells. Nucleic Acids Res. 2000;28(6):1418–23.

37. Kopera HC, Larson PA, Moldovan JB, Richardson SR, Liu Y, Moran JV. LINE-1 Cultured Cell Retrotransposition Assay. Methods Mol Biol. 2016;1400:139–56.

38. Rangwala SH, Kazazian HH, Jr. The L1 retrotransposition assay: a retrospective and toolkit. Methods. 2009;49(3):219–26.

39. Han JS, Szak ST, Boeke JD. Transcriptional disruption by the L1 retrotransposon and implications for mammalian transcriptomes. Nature. 2004;429(6989):268–74.

40. Ilik IA, Glazar P, Tse K, Brandl B, Meierhofer D, Muller FJ, et al. Autonomous transposons tune their sequences to ensure somatic suppression. Nature. 2024;626(8001):1116–24.

41. Seczynska M, Bloor S, Cuesta SM, Lehner PJ. Genome surveillance by HUSH-mediated silencing of intronless mobile elements. Nature. 2022;601(7893):440–5.

42. Kingsmore SF, Giros B, Suh D, Bieniarz M, Caron MG, Seldin MF. Glycine receptor beta-subunit gene mutation in spastic mouse associated with LINE-1 element insertion. Nat Genet. 1994;7(2):136–41.

43. An W, Dai L, Niewiadomska AM, Yetil A, O’Donnell KA, Han JS, et al. Characterization of a synthetic human LINE-1 retrotransposon ORFeus-Hs. Mobile DNA. 2011;2(1):2.

44. Richardson SR, Narvaiza I, Planegger RA, Weitzman MD, Moran JV. APOBEC3A deaminates transiently exposed single-strand DNA during LINE-1 retrotransposition. eLife. 2014;2014(3).

45. Schauer SN, Carreira PE, Shukla R, Gerhardt DJ, Gerdes P, Sanchez-Luque FJ, et al. L1 retrotransposition is a common feature of mammalian hepatocarcinogenesis. Genome Research. 2018;28(5).

46. Martin SL, Bushman D, Wang F, Li PW, Walker A, Cummiskey J, et al. A single amino acid substitution in ORF1 dramatically decreases L1 retrotransposition and provides insight into nucleic acid chaperone activity. Nucleic Acids Res. 2008;36(18):5845–54.

47. Martin SL, Cruceanu M, Branciforte D, Wai-Lun Li P, Kwok SC, Hodges RS, et al. LINE-1 retrotransposition requires the nucleic acid chaperone activity of the ORF1 protein. J Mol Biol. 2005;348(3):549–61.

48. Gerdes P, Chan D, Lundberg M, Sanchez-Luque FJ, Bodea GO, Ewing AD, et al. Locus-resolution analysis of L1 regulation and retrotransposition potential in mouse embryonic development. Genome Res. 2023;33(9):1465–81.

49. Belotserkovskii BP, Liu R, Tornaletti S, Krasilnikova MM, Mirkin SM, Hanawalt PC. Mechanisms and implications of transcription blockage by guanine-rich DNA sequences. Proc Natl Acad Sci U S A. 2010;107(29):12816–21.

50. Belotserkovskii BP, Neil AJ, Saleh SS, Shin JH, Mirkin SM, Hanawalt PC. Transcription blockage by homopurine DNA sequences: role of sequence composition and single-strand breaks. Nucleic Acids Res. 2013;41(3):1817–28.

51. Broxson C, Beckett J, Tornaletti S. Transcription arrest by a G quadruplex forming-trinucleotide repeat sequence from the human c-myb gene. Biochemistry. 2011;50(19):4162–72.

52. Beaudoin JD, Perreault JP. Exploring mRNA 3’-UTR G-quadruplexes: evidence of roles in both alternative polyadenylation and mRNA shortening. Nucleic Acids Res. 2013;41(11):5898–911.

53. Beaudoin JD, Perreault JP. 5’-UTR G-quadruplex structures acting as translational repressors. Nucleic Acids Res. 2010;38(20):7022–36.

54. Guerrero A, Herranz N, Sun B, Wagner V, Gallage S, Guiho R, et al. Cardiac glycosides are broad-spectrum senolytics. Nat Metab. 2019;1(11):1074–88.

55. Alisch RS, Garcia-Perez JL, Muotri AR, Gage FH, Moran JV. Unconventional translation of mammalian LINE-1 retrotransposons. Genes Dev. 2006;20(2):210–24.

56. An W, Han JS, Wheelan SJ, Davis ES, Coombes CE, Ye P, et al. Active retrotransposition by a synthetic L1 element in mice. Proc Natl Acad Sci U S A. 2006;103(49):18662–7.

57. Grandi FC, Rosser JM, An W. LINE-1-derived poly(A) microsatellites undergo rapid shortening and create somatic and germline mosaicism in mice. Mol Biol Evol. 2013;30(3):503–12.

58. Kannan M, Li J, Fritz SE, Husarek KE, Sanford JC, Sullivan TL, et al. Dynamic silencing of somatic L1 retrotransposon insertions reflects the developmental and cellular contexts of their genomic integration. Mobile DNA. 2017;8:8.

59. Monot C, Kuciak M, Viollet S, Mir AA, Gabus C, Darlix JL, et al. The specificity and flexibility of l1 reverse transcription priming at imperfect T-tracts. PLoS Genet. 2013;9(5):e1003499.

60. O’Donnell KA, An W, Schrum CT, Wheelan SJ, Boeke JD. Controlled insertional mutagenesis using a LINE-1 (ORFeus) gene-trap mouse model. Proc Natl Acad Sci U S A. 2013;110(29):E2706–13.

61. Wagstaff BJ, Barnerssoi M, Roy-Engel AM. Evolutionary conservation of the functional modularity of primate and murine LINE-1 elements. PLoS One. 2011;6(5):e19672.

62. Xie Y, Mates L, Ivics Z, Izsvak Z, Martin SL, An W. Cell division promotes efficient retrotransposition in a stable L1 reporter cell line. Mobile DNA. 2013;4(1):10.

63. An W, Han JS, Schrum CM, Maitra A, Koentgen F, Boeke JD. Conditional activation of a single-copy L1 transgene in mice by Cre. Genesis. 2008;46(7):373–83.

64. Newkirk SJ, Lee S, Grandi FC, Gaysinskaya V, Rosser JM, Vanden Berg N, et al. Intact piRNA pathway prevents L1 mobilization in male meiosis. Proc Natl Acad Sci U S A. 2017;114(28):E5635–E44.

65. Richardson SR, Faulkner GJ. Heritable L1 Retrotransposition Events During Development: Understanding Their Origins: Examination of heritable, endogenous L1 retrotransposition in mice opens up exciting new questions and research directions. Bioessays. 2018;40(6):e1700189.

66. Kimberland ML, Divoky V, Prchal J, Schwahn U, Berger W, Kazazian HH, Jr. Full-length human L1 insertions retain the capacity for high frequency retrotransposition in cultured cells. Hum Mol Genet. 1999;8(8):1557–60.

67. Almeida MV, Vernaz G, Putman ALK, Miska EA. Taming transposable elements in vertebrates: from epigenetic silencing to domestication. Trends Genet. 2022;38(6):529–53.

68. Lexa M, Steflova P, Martinek T, Vorlickova M, Vyskot B, Kejnovsky E. Guanine quadruplexes are formed by specific regions of human transposable elements. BMC Genomics. 2014;15:1032.

69. Sahakyan AB, Murat P, Mayer C, Balasubramanian S. G-quadruplex structures within the 3’ UTR of LINE-1 elements stimulate retrotransposition. Nat Struct Mol Biol. 2017;24(3):243–7.

70. Moran JV. Human L1 retrotransposition: insights and peculiarities learned from a cultured cell retrotransposition assay. Genetica. 1999;107(1-3):39–51.

71. Beck CR, Collier P, Macfarlane C, Malig M, Kidd JM, Eichler EE, et al. LINE-1 retrotransposition activity in human genomes. Cell. 2010;141(7):1159–70.

72. Freeman JD, Goodchild NL, Mager DL. A modified indicator gene for selection of retrotransposition events in mammalian cells. Biotechniques. 1994;17(1):46, 8-9, 52.

73. Li J, Shen H, Himmel KL, Dupuy AJ, Largaespada DA, Nakamura T, et al. Leukaemia disease genes: large-scale cloning and pathway predictions. Nat Genet. 1999;23(3):348–53.

74. Morrish TA, Gilbert N, Myers JS, Vincent BJ, Stamato TD, Taccioli GE, et al. DNA repair mediated by endonuclease-independent LINE-1 retrotransposition. Nat Genet. 2002;31(2):159–65.

75. Isomura H, Stinski MF, Kudoh A, Nakayama S, Murata T, Sato Y, et al. A cis element between the TATA Box and the transcription start site of the major immediate-early promoter of human cytomegalovirus determines efficiency of viral replication. J Virol. 2008;82(2):849–58.

